# Using the DNA language model, GROVER, to parse effects of sequence, chromatin and regulatory features on genome stability

**DOI:** 10.1101/2025.07.23.666402

**Authors:** Pierre M. Joubert, Melissa Sanabria, Anna R. Poetsch

**Affiliations:** Biomedical Genomics, Biotechnology Center, Center for Molecular and Cellular Bioengineering, Technische Universität, Dresden, Germany; Center for Advanced Systems Understanding (CASUS), Görlitz, Germany; Helmholtz Zentrum Dresden-Rossendorf (HZDR), Dresden, Germany; National Center for Tumor Diseases (NCT) partner site Dresden, German Cancer Research Center (DKFZ), Dresden, Germany; Institute for Genetics, University of Cologne, Cologne, Germany; Cologne Excellence Cluster for Cellular Stress Responses in Aging-Associated Diseases (CECAD), University of Cologne, Cologne, Germany; Institute for Genome Stability in Aging and Disease, University of Cologne, Cologne, Germany

**Keywords:** genome stability, deep learning, large language model, interpretable machine learning, epigenetics

## Abstract

**Abstract:** Genome stability is shaped by DNA sequence and chromatin context, but their relative contributions to double-strand break (DSB) sensitivity remain unclear. We show that the DNA language model, GROVER, can infer DSB location based on sequence. DSB hotspots tend to contain GC-rich sequences that belong to promoters, genes and short interspersed nuclear elements (SINEs). Additionally, we identified several specific short sequences (tokens) that are associated with modulating DSB sensitivity. Another model using chromatin and genome regulatory features outperforms the sequence-only model, highlighting complementary and cell-type specific information. Integrating sequence and genome biological features yields the best performance, demonstrating their synergy. Analyzing this model revealed that, dependent on the sample, genome stability information encoded in H3K36me3 and DNase-seq can be learned from the sequence, but not H3K27ac or H3K9me3. Embedding chromatin data directly into the GROVER architecture enabled cell-type specific modeling with performance matching the full chromatin feature model. Our results suggest that while chromatin and regulatory context provides important information, such as cell-type specificity, much of the information shaping DSB patterns is already encoded in the DNA sequence itself. Our integrative modeling approach not only reveals DSB patterns but also provides a generalizable strategy for tracing predictions in genomic data.

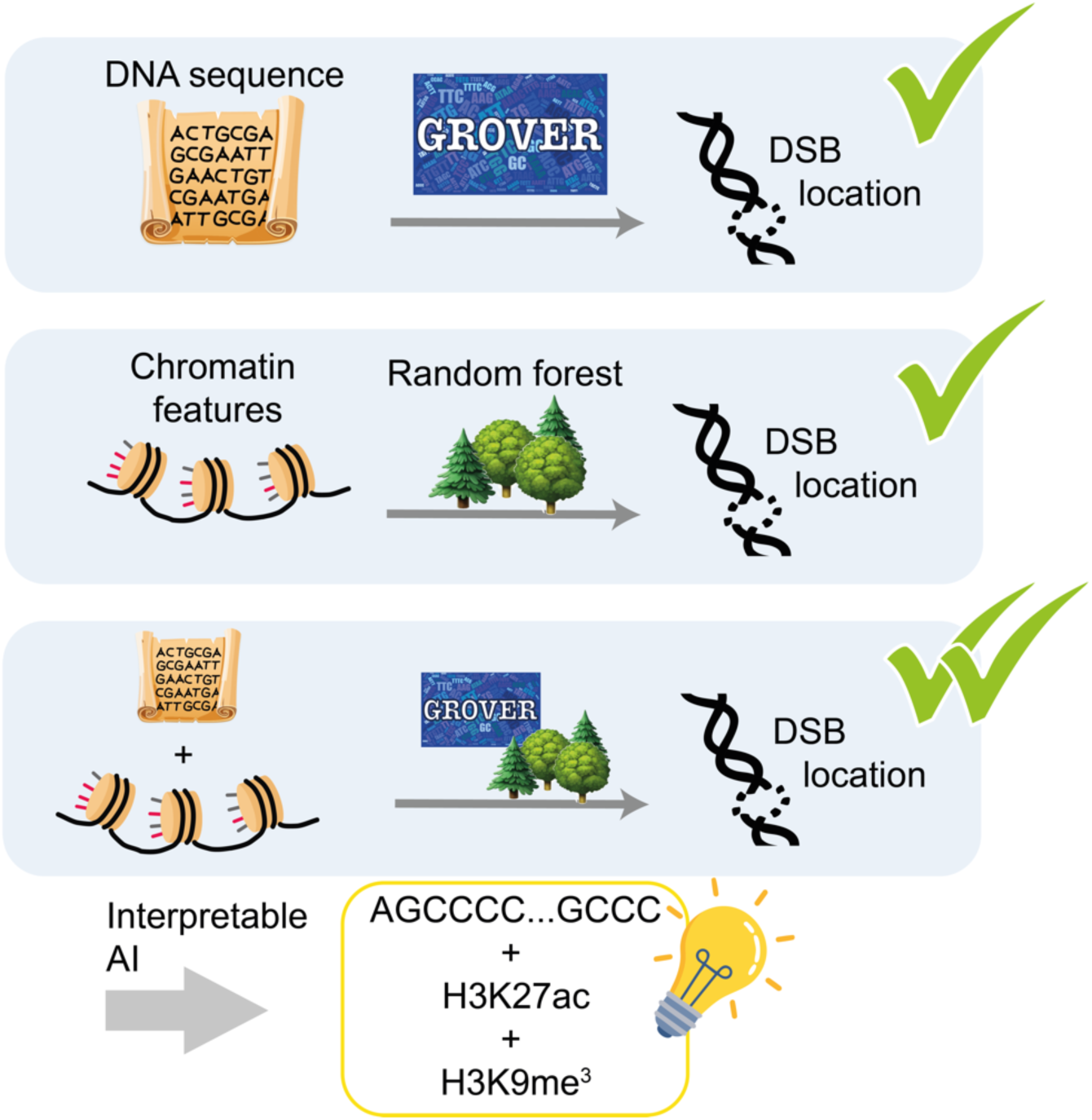

## Background

Organisms must protect their genomes to function properly. Genome instability can lead to disease but also serves as a driver of evolution. Although it is well established that chromatin features and DNA sequence context influence genome stability, the extent and the precise mechanisms remain unclear. One way to study genome stability is by mapping the locations of double-strand breaks (DSBs). Previous studies have established that DSBs are not randomly distributed across the genome but instead show enrichment or depletion in specific chromatin and sequence contexts. For example, DSBs are frequently associated with genes, and especially promoters, which is attributed to topoisomerase activity [1] and R-loop formation [2,3], which often occurs at G-quadruplexes [4–6]. This association, along with the co-localization of CpG islands and promoters [7,8] and the presence of high levels of DSBs at telomeric repeats [9], results in a correlation between DSBs and GC content. Similarly, Alu elements tend to be found in GC-rich regions [8] and are known to be a common repair substrate for homology-directed repair [10,11]. Other repeats, such as Long Interspersed Nuclear Elements (LINE) [12], microsatellites [13,14], and palindromic AT-rich repeats [15] are also known as hotspots for DSBs. Common fragile sites, which are often found in long, AT-rich genes, are also thought to be enriched in DSBs [16]. Finally, CRISPR-induced DSBs have been shown to exhibit sequence-dependent repair [17–19]. Taken together, these studies indicate a strong sequence-specificity in DSB locations and repair. However, it is unclear to which extent the sequence alone is sufficient to understand and model the DSB landscape and which additional information from genome biological features and tissue identity is necessary.

Finding patterns in genome biological features that define where DSBs occur is complicated by the fact that sequence elements, like genes and repeats, have strong trends in which chromatin context they are found. For example, active transcription, which occurs within genes, is associated with DSBs [20]. Active transcription is also associated with histone marks that themselves correlate with DSBs. These marks include the active enhancer and promoter mark H3K27ac [21–24] and the active promoter mark H3K4me3 [21,24,25]. Similarly, H3K36me3, which marks actively transcribed gene bodies, plays a significant role in coordinating DSB repair [26]. H3K9me3 [27] and H3K27me3 [28–30], which mark constitutive and facultative heterochromatin, respectively, both play a role in DSB repair [27,31,32]. Additionally, CCCTC-binding factor (CTCF) is of particular interest, as it plays a direct role in DSB repair [33,34] and chromosome loop anchors bound by CTCF are prone to DSBs [35]. Replication timing plays a crucial role in DSB generation as well, both at late-replicating common fragile sites [36,37] and in early-replicating Poly(dA:dT) sites [38,39] and promoters [1–3]. Furthermore, recently developed DSB-sequencing technologies such as DSBCapture [40], BLISS [41], and its variant sBLISS [42] have enabled the rapid and straightforward genome-wide quantification of DSBs *in situ*. These methods provide a snapshot of DSB locations, which are shaped by repair pathway decisions and differences in repair speed. These methods have enabled characterization of the overall, genome-wide association of DSBs with chromatin accessibility [40,43,44], CTCF-binding [40,43,44] and early replication [43].

Many genome biological features can themselves be inferred from DNA sequence with good performance [71] and are thus to some extent by themselves encoded, which makes it difficult to disentangle the relative contributions. CRISPR-based experiments using the same sequence inserted into different contexts have shown that chromatin profiles influence the pathway decisions of DSB repair in a sequence-agnostic way [45–47]. However, understanding how DSB sensitivity changes in different chromatin and regulatory contexts is a difficult task given the complexity of the underlying sequence. Therefore, powerful tools that can capture the non-linear interactions between features within these sequences are required.

Recent advancements in deep learning have enabled the development of DNA language models, which can interpret these complex interactions within the sequence and can themselves be highly interpretable thanks to their self-attention mechanism [48]. In practice, however, interpreting these models in a biomedical context is difficult [49–51], and special care must be given to choices like model architecture and vocabulary choice to enable these models to understand sequence context [52]. We have shown that our model, GROVER (Genome Rules Obtained Via Extracted Representations), learns sequence context and many features of DNA without supervision, including GC content, genes, repetitive elements and replication timing [53]. GROVER can also be fine-tuned on additional data to perform many tasks, including predicting CTCF-binding and recognizing promoters. These findings make GROVER particularly well suited for understanding the relationship between the genome, the chromatin and regulatory state and the DSB landscape.

DSB locations are shaped by both the DNA sequence and genome biological features, with the chromatin and regulatory state itself being influenced in part by the underlying sequence (**Fig. 1A**). In this study, we aim to disentangle these intertwined contributions and assess the extent to which each shapes the DSB landscape using interpretable machine learning approaches. We show that GROVER can be fine-tuned to predict DSB sensitivity, indicating that DSBs can be modeled using the sequence alone. However, combining GROVER with genome biological features results in a better-performing model, which demonstrates that these genome biological features shape the DSB landscape independent of the sequence. We also show, using interpretable machine learning techniques, that certain genome biological features overlap with what can be learned from the sequence, and others do not, indicating that these features could be important for determining the cell type-specificity of DSB locations. By integrating these non-redundant features into the GROVER framework, we learn about the interaction of the DNA sequence with the chromatin and regulatory state for genome instability. Such an approach is generally suited to disentangle the contributions of sequence and chromatin context to genome biology.

**Fig. 1.**
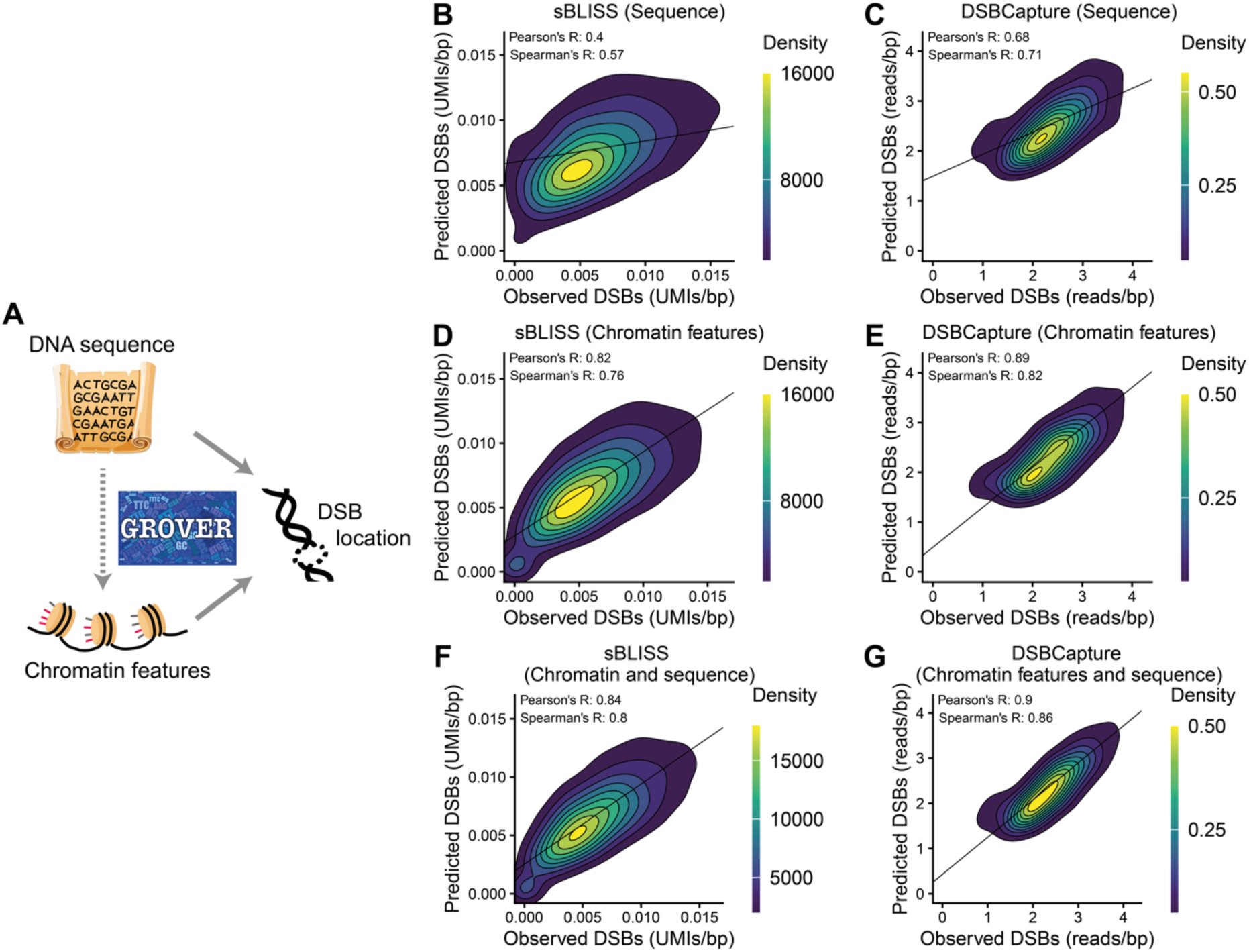
Sequence and chromatin context both contain information that can help model double strand breaks (DSBs). **A**. DSB locations are shaped by both the DNA sequence and the chromatin context, with the chromatin state itself being influenced in part by the underlying sequence. DNA language models like GROVER, combined with interpretable machine learning approaches, can be used to disentangle these interactions. **B and C.** Performance of GROVER model fine-tuned to predict sBLISS data (**B**) and DSBCapture data (**C**) using the DNA sequence alone. **D and E.** Performance of random forest model trained on chromatin features to predict sBLISS data (**D**) and DSBCapture data (**E**). **F and G.** Performance of random forest model trained on chromatin features combined with the predictions of the GROVER fine-tuned model to predict sBLISS data (**F**) and DSBCapture data (**G**). DSB data is shown as molecular identifier (UMI) counts (sBLISS) or read counts (DSBCapture) averaged per base pair over the length of each window.

## Results

### The DNA sequence and the genome biological state provide distinct information that can be used to model double strand breaks

To study the relationship between the DNA sequence, the genome biological context and genome stability, we re-analyzed two previously published DSB sequencing datasets, one using sBLISS in breast cancer MCF7 cells [43] and one using DSBCapture in epidermal keratinocyte NHEK cells [40]. While both technologies involve a blunting step to prepare DNA ends for adapter ligation, a key difference is that sBLISS incorporates unique molecular identifiers (UMIs) and uses T7-based *in vitro* transcription for linear amplification, enabling quantitative measurement of DSBs [41,42]. In contrast, DSBCapture relies solely on PCR amplification [40]. In our dataset, this likely results in DSBCapture being more sensitive to rare DSBs but prone to overestimating the frequency of recurrent breaks due to PCR duplicates, as well as introducing sequence-dependent amplification biases, which may distort the observed distribution of DSBs [54,55] and impair comparison of absolute DSB counts between samples. To avoid potential biases in DSB locations caused by DSB-inducing treatments [56], both datasets we used were generated from cells that were not treated and have no reported clear DNA repair deficiency.

To determine whether DSB locations can be inferred using just the genome, we fine-tuned GROVER for this task. We computed the DSB counts from our datasets for sequence windows that are of length equal to the maximum input size of the model (2,075 base pairs on average). GROVER predicts DSB counts generated from sBLISS and DSBCapture datasets with a Spearman’s R of 0.57 and 0.71, respectively (**Fig. 1B and 1C**), which shows that DSBs can be modeled using the DNA sequence data alone, yet with suboptimal performance.

In parallel, we trained a model only on genome biological features, which represent both context dependency, e.g. tissue identity, and also correlative sequence characteristics with the stability measure. Following previously published methods [43], we gathered various datasets from ENCODE [57] and used this data to train a random forest for prediction of DSB counts on the same windows as we used for GROVER and compared these also to other machine learning architectures. Despite the small window sizes, the models are still able to predict DSB counts generated by both datasets, with a Spearman’s R for the random forest of 0.76 for the sBLISS data and 0.82 for the DSBCapture data and equivalent performance for other architectures (**Fig. 1D and 1E Additional file 1: Fig.S6-10**). This shows that the chromatin and regulatory state is more predictive of the DSB sequencing data than the model that is based on genome sequence alone, it is however unclear how much the shared sequence characteristics between the features and the DNA damage output contributes to the predictions.

To understand the redundancy or potential synergy between sequence and genome biological context, we combined the models by using the fine-tuned GROVER model’s predictions as a feature of the models trained on genome biological features (**Fig. 1F and 1C, Additional file 1: FigS6-10)**. The combined random forrest model performs slightly better than either of the previous models, with a Spearman’s R of 0.80 for the sBLISS data and 0.86 for the DSBCapture data with equivalent performances for the other architectures. The increase in the models’ performance compared to the biological feature model indicates that GROVER learns some additional information from the sequences that cannot be learned from the shared sequence characteristics of the genome biological context. To quantify this effect, we measured the difference in R^2^ values between the biological feature random forest model and the combined model and found that GROVER predictions explain an additional 3.5% of the variance for the sBLISS data and 2% of the variance for the DSBCapture data (**Additional file 1: Fig. S1A and S1B**) that is not encoded in the genome biological features. Similarly, we found that GROVER predictions are correlated with the residuals from the genome biological feature model with a Spearman’s R of 0.17 for the sBLISS data and 0.20 for the DSBCapture data (**Additional file 1: Fig. S1C and S1D)**.

In summary, these results indicate that the genome sequence and the regulatory or chromatin context contain both overlapping and distinct information that can be used to predict DSBs. We therefore sought to understand where these overlaps and distinctions might be by further analyzing the features and the models that we trained them on.

### DNA sequences display features that are associated with double strand breaks

To better understand what information about DSBs can be encoded within the genome directly, we explored sequence-encoded features that are associated with DSBs. As expected, we found that GC content is correlated with DSBs, with a Spearman’s R of 0.23 for the sBLISS data and 0.62 for the DSBCapture data (**Fig. 2A and 2B**), which is unlikely to be purely of technical origin [40]. These correlation coefficients are meaningfully lower than GROVER’s performance, indicating that the sequence contribution to DSB counts is greater and more complex than just its GC content alone. Additionally, we found that the presence of a promoter in a sequence results in a doubling of DSBs on average (**Fig. 2C and 2D**), in line with previously published results [1–3]. Even though promoters often have high GC content, this association does not appear to be driven purely by GC content (**Additional file 1: Fig. S2A and S2B)**. The presence of short interspersed nuclear elements (SINEs), including Alus, and genes are also associated with increased DSBs, which is expected given that these three features are correlated with the presence of promoters [8]. These results hint that DSB hotspots are characterized by specific patterns within the sequence itself. While such complex patterns can be difficult to learn using conventional machine learning models, DNA language models such as GROVER can learn their representations even during pre-training [53]. This likely explains why GROVER can model DSB locations using the sequence alone.

**Fig. 2.**
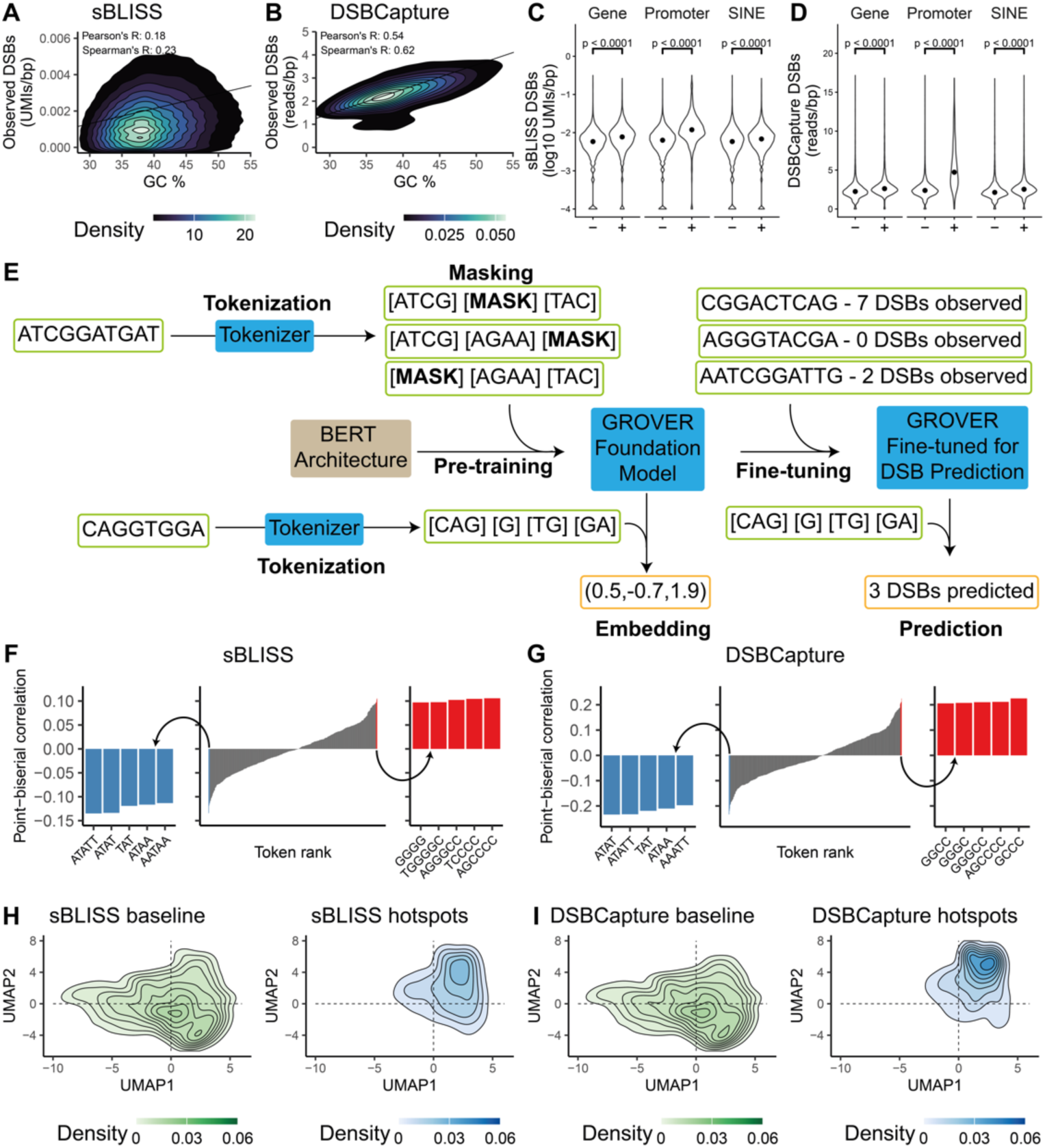
Sequence features are associated with double strand breaks (DSBs). **A and B.** Correlation between sequence GC content and sBLISS data (**A**) or DSBCapture data (**B**). **C and D.** Comparison of sBLISS data (**C**) and DSBCapture data (**D**) across sequences categorized by the presence of a gene, promoter, or short interspersed nuclear element (SINE) (p < 0.0001 by Wilcoxon rank test). Points represent median values. **E.** Explanation of GROVER model data processing, pre-training and fine-tuning workflow (see main text). For the sake of visual clarity, a few simplifications have been made. The embedding output from the pre-trained model has 768 dimensions, but only three are shown here. Sequences input into the model are 510 tokens long (or 2,075 base pairs on average), but only a few tokens are shown here. Additionally, DSB counts are normalized for sequence window length in our data, which is not shown here. **F and G.** Point-biserial correlation between the presence of each token within the GROVER vocabulary and sBLISS data (**F**, UMIs per base pair in each window) or DSBCapture data (**G**, reads per base pair in each window). **H and I.** Uniform manifold approximation and projection (UMAP) analysis of pre-trained GROVER embeddings separated by DSB level, according to sBLISS data (**H**) and DSBCapture data (**I**). Sequences are separated based on whether they are in the top 10% (hotspots) or bottom 90% (baseline) of sequences.

Before a sequence can be used as an input to GROVER for fine-tuning, it must be separated into tokens using the model’s tokenizer (**Fig. 2E**). These tokens are of variable lengths, which results in variable lengths of the input sequences and requires the DSB counts to be normalized by sequence length in our dataset. An enrichment of specific GROVER token types within high DSB sequences could indicate a pattern of enrichment of specific short sequences in areas of the genome that frequently experience DSBs. When we analyzed these enrichments, we found that, across both datasets, GC-rich tokens like “AGCCC” are frequently associated with increased DSBs while AT-rich tokens like “ATATT” are anticorrelated (**Fig. 2F and 2G**).

During pre-training, GROVER learns patterns in sequences in a self-supervised fashion, i.e., without the addition of any additional biological information [53]. Using a DNA sequence as input for GROVER without fine-tuning the model results in a vector of numbers, known as an embedding, which can be analyzed to interpret what patterns exist within the sequence (**Fig. 2E**). Generating a uniform manifold approximation and projection (UMAP) of these embeddings, we observed separation between sequences with low and high DSBs, which indicates that regions of the genome that are particularly prone to instability can be differentiated by their sequence alone (**Fig. 2H and 2I**). This analysis also shows that sequences containing similar features are DSB hotspots in both DSBCapture and sBLISS datasets, despite the differences in methods and cell types. Overall, our results support the presence of patterns within the sequence itself that are reflected in the associated location of DSBs.

### Patterns within genome biological features are associated with double strand breaks

To understand what patterns within the chromatin and regulatory context might be associated with DSBs, we calculated the correlation between the genome biological features and DSB sequencing data. We found that sBLISS data is correlated with all chromatin features, including counter-intuitively also repressive histone marks like H3K9me3, yet to a larger extent H3K36me3 and H3K27ac histone marks as well as RNA polymerase 2 (Pol2) pausing and RepliSeq data (Spearman’s R of 0.43, 0.42, 0.37 and 0.34, respectively, **Fig. 3A**), while the DSBCapture data is correlated with RepliSeq, DNase-Seq and Pol2 pausing data (Spearman’s R of 0.65, 0.64 and 0.51, respectively, **Fig. 3B**). Despite using smaller sequence windows, our results align with previously published results [40,43]. The generally positive correlation of histone marks with sBLISS may hint towards ChiP-Seq sequencing artefacts of preferential sequencing in accessible DNA, which are addressed in more detail below. In addition, the differences between the two datasets may be caused by the specifics of the sequencing technologies used. However, given the fact that these details should largely only affect PCR duplicates, it is more likely that the differences in the chromatin features, in particular the differences for heterochromatic marks, can be attributed to variable DSB and chromatin profiles of the two cell lines, or variation in ChIP-seq data quality.

**Fig. 3.**
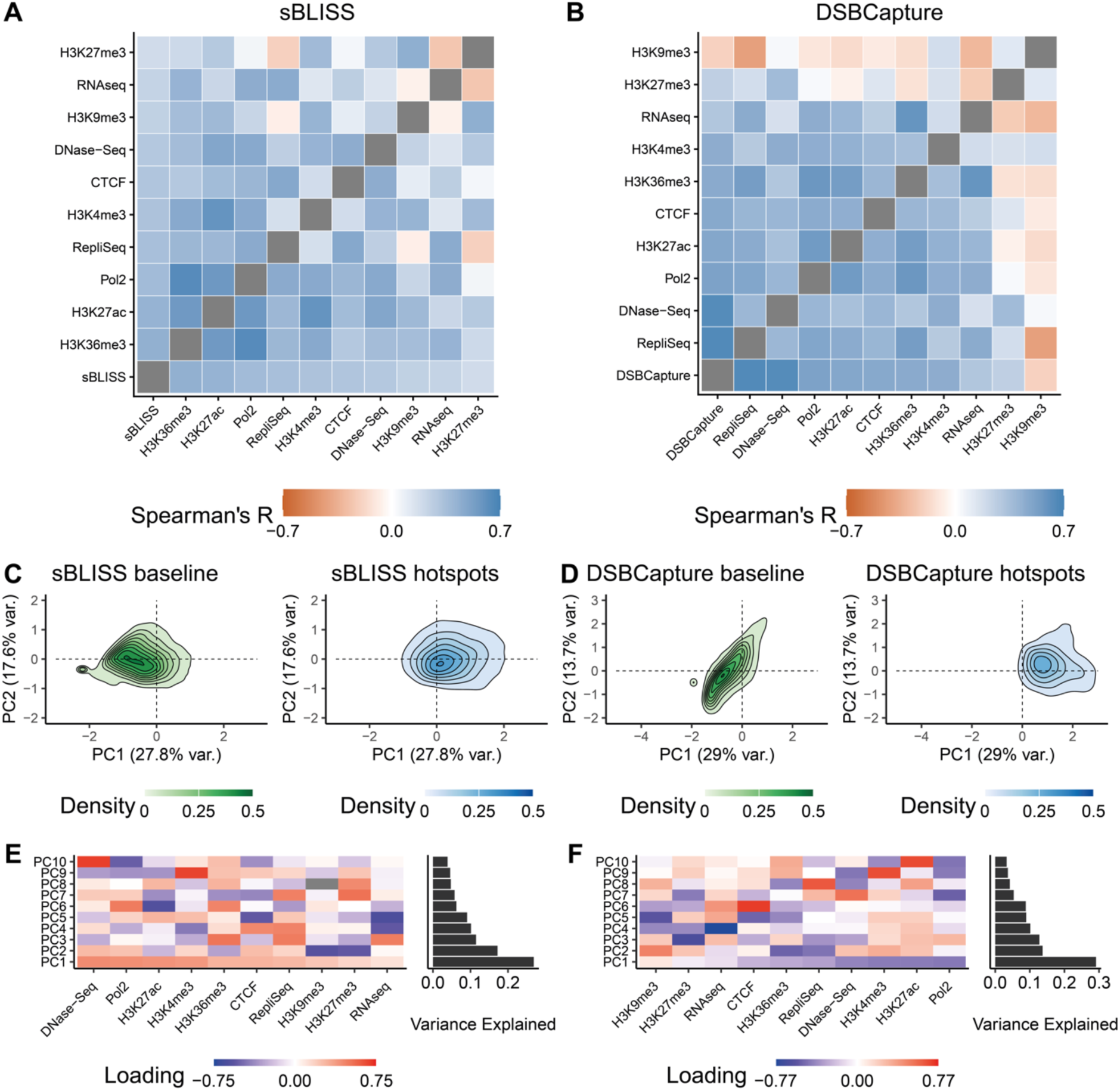
Chromatin features are correlated with double strand breaks (DSBs). **A.** Spearman correlation between MCF7 chromatin features and sBLISS data (**A**, unique molecular identifiers per base pair) or DSBCapture data (**B**, reads per base pair), ranked by their correlation with the DNA double strand break measurement. **C and D.** Principal components analysis (PCA) of MCF7 (**C**) and NHEK (**D**) chromatin features. Sequences are separated based on whether they are in the top 10% (hotspots) or bottom 90% (baseline) of sequences. **E and F.** Loading of each MCF7 (**E**) and NHEK (**F**) chromatin feature within each PC, as well as the variance explained by each PC, ranked by the loadings of PC1.

We then asked whether the variation in genome biological features across the genome could separate sequences experiencing different numbers of DSBs. We performed a principal component analysis (PCA) and found that sequences with high DSB levels are linearly separated from others along the first principal component (PC1), which explains more than 25% of the variance in the data (**Fig. 3C and 3E**). The correlation between PC1 and DSB levels suggests that some of the dominant patterns in genome biological variation are aligned with DSB locations. PC loadings show that PC1 for both datasets is primarily driven by DNase-seq, RNA Pol2, H3K27ac and H3K4me3, indicating that a sizeable part of the variance in these datasets can be attributed to the same genome biological features, though their relative contributions differ (**Fig. 3D and 3F**). Additionally, we used UMAP to visualize the chromatin and regulatory data in non-linear space and again observed separation between low and high DSB sequences (**Additional file 1: Fig. S3**). These results confirm previous findings that certain chromatin features align well with DSBs and that these features can be used to characterize DSB hotspots.

With the aim of understanding which genome biological features have a central role in shaping the DSB landscape in the genome, we assessed the relative importance of these features within our random forest models by calculating the decrease in the mean-squared error (MSE) of the model when permuting each feature in the training data. We found that H3K36me3, H3K9me3, and H3K27ac are the most important features for the sBLISS data (**Fig. 4A**) with H3K9me3 and H3K27ac swapping their relative order with different machine learning architectures (**Additional file 1: FigS6-8 E**). DNase-seq is the most important for the DSBCapture model by far, followed by H3K27ac and RepliSeq (**Fig. 4B**) with variable order dependent on the machine learning architecture (**Additional file 1: FigS6-8 M**). While the importances we observed reflect the ones published by Ballinger *et al*. [43], one major difference is the smaller importance of RepliSeq in our models. We observed the same results when we repeated our analysis using analogous 50,000 base pair windows (**Additional file 1: Fig. S4A-D**).

**Fig. 4.**
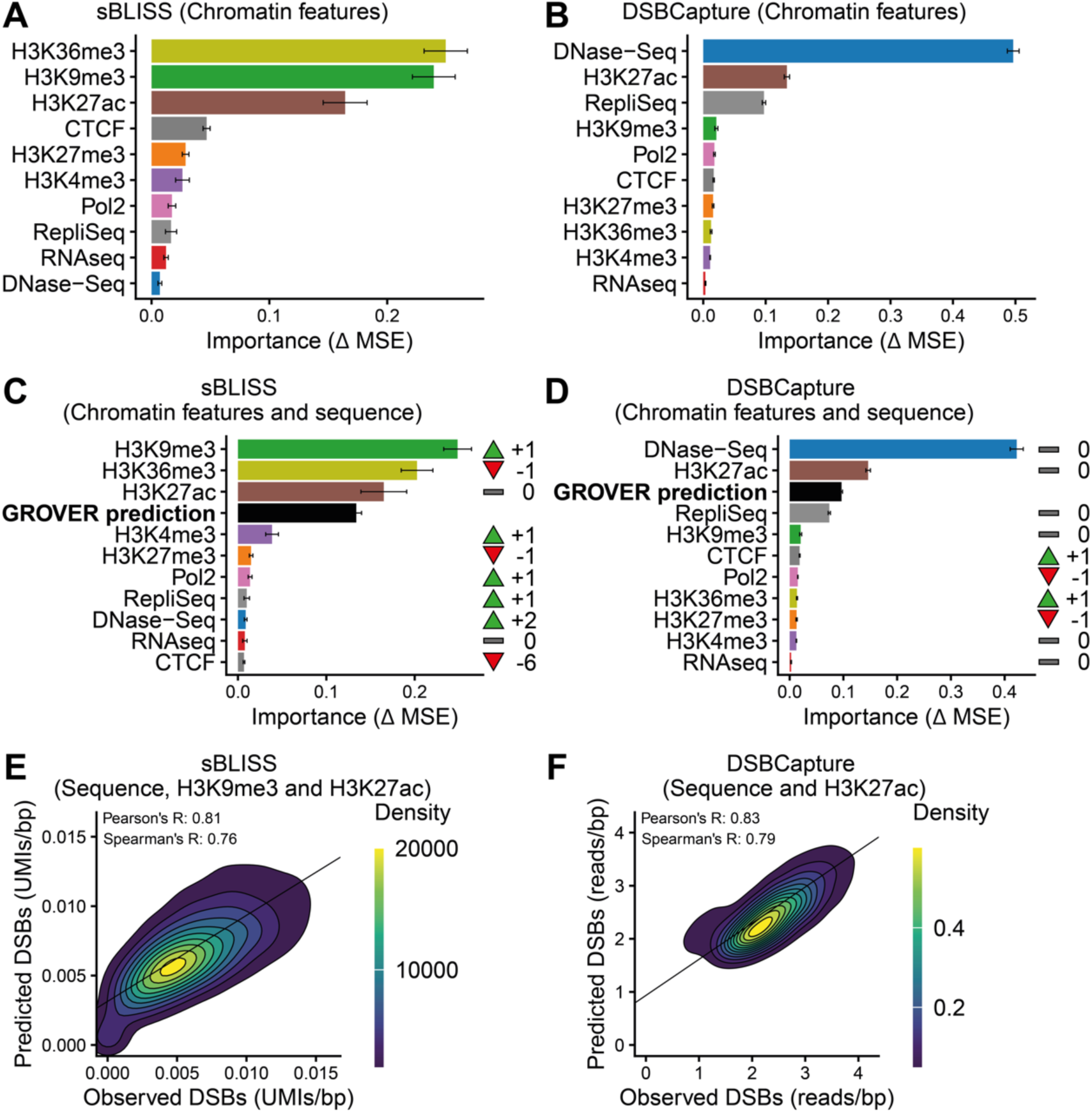
Sequence context is an important feature for predicting double strand breaks (DSBs). **A** and **B.** Importances of features used by the random forest model trained on chromatin features to predict sBLISS data (**A**) and DSBCapture data (**B**). **C and D.** Importances of features used by the random forest model trained on chromatin features combined with the predictions of the fine-tuned GROVER model to predict sBLISS data (**C**) and DSBCapture data (**D**). Arrows represent changes in rank of feature importances compared to panel **A** and **B**, respectively. **E and F.** Performance of GROVER model fine-tuned on DNA sequence and limited chromatin features to predict sBLISS data (**E**) and DSBCapture data (**F**). Error bars represent standard deviation from the mean. Importances are measured using the change in mean squared error (MSE) of the model when a feature is shufled in the test data. DSB data is shown as molecular identifier (UMI) counts (sBLISS) or read counts (DSBCapture) averaged per base pair over the length of each window.

Though we did not include this feature in our random forest model because it is not expressly biologically relevant, while quality checking our chromatin mark and regulatory feature data, we noticed that the corresponding ChIP-Seq input datasets are correlated with both sBLISS and DSBCapture data (Spearman’s R of 0.45 and 0.27, respectively, **Additional file 1: Fig. S5A- B**). When we added this data to new machine learning models, it became the top feature in the sBLISS random forest model, dwarfing all other features and substantially boosting the performance of the model, yet less so for DSBCapture (**Additional file 1: Fig. S4E-H, S5C-F**). Also in the other machine learning models, input is recognized as a major feature for sBLISS in MCF7, but not for DSBCapture in NHEK cells (**Additional file 1: FigS6-8 G**). Assuming there is no major sequencing bias in sBLISS, which is supported by the original publication [41], this result could suggest overlap in sequences that are prone to DSBs and sequences that are prone to being over-sequenced in a ChIP-Seq input sample due to a bias in chromatin accessibility, associated fragmentation and subsequent size selection. Therefore, the input may already represent DNA accessibility, which is a feature with strong correlation with DSBs.

In summary, our analysis of these genome biological features and our random forest models demonstrates that patterns in these features are associated with DSBs and that they can be used to model DSBs.

### The chromatin and regulatory context and the genome contain non-overlapping patterns associated with double strand breaks

After identifying patterns within the sequence and the chromatin and regulatory context that are associated with DSBs, we sought to compare these patterns to determine their similarities and differences. When we added GROVER’s predictions to our machine learning models that previously only contained genome biological features, we found that the GROVER prediction is the top 4 most important features for the sBLISS random forest model and XGBoost, the top 3 for MLP and LightGBM, and for DSBCapture the top 3 for the random forest and top 2 for the other model architectures (**Fig. 4C and 4D, Additional file 1: FigS6-8 F,N**). This indicates that GROVER learns important DSB information from the sequence that is not present in the chromatin mark and regulatory feature datasets. This is expected given the increase in performance when GROVER is included in the genome biological feature model (**Fig. 1F-G, Additional file 1: FigS6-8 B,J,S10**). To confirm GROVER’s importance in the combined random forest model, we also performed a complementary approach to measure feature importance. Rather than permuting individual features, we also removed each feature from the dataset one at a time and measured the decrease in MSE with the reduced feature set. With this approach we found that GROVER is the most important feature for the sBLISS data and the second most important feature for the DSBCapture data (**Additional file 1: Fig. S1E-F**).

In this type of importance analyses, features encoding similar information tend to “steal” importance from each other [58], so analyzing how these permutation-based importances change when GROVER and therefore sequence information is explicitely included in the model could give us a better idea of how DSB sensitivity is encoded at the genomic level and in the genome biological features. For the sBLISS data and random forest model, we observed that H3K36me3, H3K27me3, and especially CTCF decrease in importance, indicating that the information learned by GROVER in the sequence is partially redundant with the information encoded in these features, or completely redundant, in the case of CTCF (**Fig. 4A and 4C**), yet the relative feature importance of histone marks also varies by machine learning architecture (**Additional file 1: FigS6-8 F**). CTCF binding can be predicted by GROVER very well [53], which suggests that it learned the link between CTCF binding and DSBs [35,59], from the sequence alone. We also found that H3K9me3, H3K4me3, and H3K27ac retain similar importance when GROVER predictions are included, hinting that these chromatin marks contain patterns associated with DSBs that cannot be learned from the sequence alone.

For the DSBCapture data, the ranks of the genome biological features remain largely the same after the inclusion of the GROVER predictions, though DNase-seq and RepliSeq lose some of their importance, which indicates that some of the information contained within these features overlaps with the patterns found within the sequence (**Fig. 4B and 4D, Additional file 1: FigS6-8 N**). These results are likely influenced by the dominance of DNase-seq as a feature in the DSBCapture models. However, we observed once again that the importance of H3K27ac does not change for this dataset, which supports that this chromatin mark encodes information that we cannot learn from the sequence.

To provide GROVER with complementary chromatin feature information, we incorporated the sequence-complementary chromatin marks directly within the architecture of the GROVER model. We used H3K9me3 and H3K27ac data for the sBLISS model and H3K27ac data for the DSBCapture model and improved performance over the original fine-tuned GROVER models for both sBLISS and DSBCapture, with a Spearman’s R of 0.76 and 0.79, respectively (**Fig. 4E- F**). For the sBLISS data, the fine-tuned GROVER model even matches the performance of the genome biological feature model (**Fig. 1D**). This indicates that the DNA sequence combined with two minimal chromatin features that represent the cell-type information are sufficient to model DSBs as well as the full panel of genome biological features.

## Discussion

In this study, we sought to disentangle and understand the influences of the chromatin and regulatory context and the DNA sequence on genome stability. To this end, we successfully fine-tuned GROVER to predict DSBs using only DNA sequences. This result shows that DSBs can be modeled directly using the genome, and our analysis of sequence-encoded features shows how these features appear to influence DSB sensitivity. To identify whether the sequence alone is enough to fully model DSBs, we compared our fine-tuned GROVER model’s performance to the performance of genome biological feature-only models. We found that GROVER performs worse than these models, showing that the genome biological context contains additional useful information for modeling DSBs that cannot be learned from the sequence. Besides technical reasons, this difference in performance can be explained by additional information on cell-type that cannot be interpreted from the DNA sequence alone. We also found that the two datasets differ in which features are particularly predictive of DSBs. This may be due to differences in sequencing technologies or biological differences between cell types. The minor differences between the two technologies point towards the latter, though this is difficult to fully answer given the current availability of datasets. Biological differences could originate from the fact that MCF7 is a cancer cell line while NHEK is not. For example, homologous recombination is reported to be upregulated in MCF7 cells [60]. The differences we observed between the two datasets highlights the need for more high-quality DSB-sequencing datasets in untreated cells. Additionally, technologies such as DSBCapture and sBLISS provide a snapshot of DSBs and their repair, and it is challenging to differentiate the two and interpret the associated patterns accordingly. Selective DNA repair rather than DNA damage impact could explain why H3K36me3 is such a strong predictor for the sBLISS data, for example, given the histone mark’s important role in DNA repair [26]. While the genome biological features we identified largely reflect the ones published by Ballinger *et al*. [43], one major difference is the reduced importance of RepliSeq in our models. While we used data from the same published datasets as Ballinger *et al*., our analysis used the Wavelet-smooth Signal tracks rather than the Summed Densities tracks from ENCODE. The description of these file types as well as the original RepliSeq publication support our choice [61,62]. The importance of the ChIP-Seq input data in our models that included this feature could explain why Ballinger *et al*. found that RepliSeq is such an important feature while we did not.

When we combined the modalities, we found that the combined models’ performances are greater than either of their parts, indicating that the sequence and chromatin state provide synergistic information. Furthermore, we found that the sequence itself, in the form of GROVER’s predictions, is one of the most important features of this combined models. These results show that the genome and the chromatin context capture both shared and distinct information. Analyzing the importances of this combined model and how they differ from the genome biological feature-only model revealed that some of the DSB information provided by chromatin features like H3K36me3 can be directly learned from the sequence, which suggests that the sequence is related to the location of these features. This analysis also shows that certain features like H3K27ac do not overlap with what GROVER learns from the sequence, hinting that these features are not influenced by the sequence in the context of DSBs. Broadly, these results underscore the importance of the chromatin state in shaping the DSB landscape, while also further raising the question of which chromatin features act directly or merely reflect underlying DNA sequence patterns.

Our findings reveal that the sequence alone is not sufficient to fully model DSBs. However, when we added the sequence-independent chromatin features of just one or two histone marks directly to the GROVER architecture and fine-tuned the model, we found that the model’s performance is competitive with the chromatin feature model. Interpreting the importances of machine learning models is a powerful tool to identify the genome-wide relevance of these features. However, sophisticated approaches for the interpretation of DNA language models, such as attention analysis, have been developed [49–51] that could allow the fine-grained interpretation of these patterns and identify in which sequence contexts particular chromatin features are linked to DSB location. We found that specific tokens are associated with DSBs and confirmed previously published results regarding the association of high GC content and promoters with DSBs [1–5,7–9]. Several known mechanisms lead to accumulation of DSBs in association with GC rich DNA, e.g. the accessibility of DNA during regulatory activities in promoters, topoisomerase activity, and processes like transcription itself [72]. Likewise, G-rich sequences are on the one hand particularly prone to oxidation [63], but on the other hand reduced in 8-oxoG/ apurinic sites [68–70]. However, exposure to oxidizing agents is cell-type specific as well as gene regulation and gene activity. Therefore, multiple mechanisms link DSB accumulation both to the DNA sequence and its regulatory chromatin context. In addition to the damage accumulation, also DSB repair is widely dependent on chromatin context [73], where chromatin marks affect pathway choice via impacting the proteins involved to a variable extent. The promoter and enhancer accessibility marks H3K27ac and DNase-sensitivity, favor N-synergy [Verarga], an association with Non-homologous end joining (NHEJ). They are the most sequence-complementary features in our study and still are both associated with GC rich DNA. Yet this ostensible contradiction can be explained via the tissue dependency of the sequence-encoded potential for accessibility. Therefore, not also the generation of DSBs, but also local differences of DNA repair can be both sequence and genome biological context dependent. It remains to be investigated how much the regional differences are due to DSB accumulation or its repair, and how much the sequence versus the chromatin and regulatory state contribute to each.

Interpretable machine learning provides an attractive avenue. However, the associated interpretability analyses are still maturing, and their use in a biomedical context remains challenging [49–51]. Because the chromatin features we integrated within GROVER appear to be sequence-independent, upscaling such an approach may allow the associated models to interpret the sequence in a cell-type specific manner. In addition, this strategy of multimodal learning improves performance, which is often associated with reduced noise and greater interpretability, without introducing redundant features that overlap much with the sequence. As interpretation methods continue to develop, incorporating a minimal set of chromatin features may offer a practical way to include cell-type specific information while preserving model clarity.

## Conclusions

Our study advances the understanding of how both DNA sequence and genome biological context contribute to genome stability. By leveraging interpretable machine learning approaches, we demonstrate that while DNA language models like GROVER can capture important sequence-level information related to DSBs, the chromatin and regulatory state provides synergistic information. Importantly, we show that incorporating a minimal set of sequence-independent chromatin features can substantially improve performance without sacrificing interpretability, offering a promising strategy for building biologically meaningful models. This work highlights the potential of combining deep learning with genomic and chromatin data to understand the relationship between DNA sequence and genome biology. We hope these insights will inspire future efforts to use DNA language models not just for prediction but for revealing the principles by which sequence and chromatin context shape the genome’s stability and function across diverse biological settings.

## Methods

Machine learning and data preparation scripts were written in Python (v3.11.5) with scikit-learn (v1.2.2), numpy (1.26.2), pytorch (v2.7.1+cu126), pandas (v2.1.1), and pyranges (v0.1.2). Plotting and statistical testing scrips were written in R (v4.4.0) with ggplot2 (v3.5.1), dplyr (v1.1.4), readr (v2.1.5), scales (v1.3.0), viridis (v0.6.5), reshape2 (v1.4.4), tidyverse (v2.0.0), ggrastr (v1.0.2), ggsignif (v0.6.4), RColorBrewer (v1.1.3), patchwork (v1.3.0) and reticulate (v1.40.0).

### Double strand break data

The previously published, GROVER-tokenized *Homo sapiens* (human) hg19 (GRCh37) genome was divided into 510 token windows [53]. Any windows containing Ns were removed from the dataset. DSBs counts were then determined according to previously published methods [40,41,43] and by adapting the code obtained from Ballinger et al. [43]. For the sBLISS data, FASTQ files were downloaded from the Sequence Read Archive (SRP150602) and filtered to verify that each read contained the expected 8 bp UMI and the 8 bp sample barcode while allowing one mismatch per barcode using *scan_for_matches* (https://blog.theseed.org/servers/2010/07/scan-for-matches.html). Filtered reads were then mapped to the hg19 genome using BWA-MEM (v0.7.17-r1198-dirty) [64] and alignments with a mapping quality lower than 30 were removed using SAMtools (v1.3.1) [65]. Read alignments were used along with their UMIs to produce a strand-aware, deduplicated list of breakpoint locations and counts as a bedGraph file which was then filtered for PCR duplicates using umi_filter.py [43] before it was converted to a bigWig file using bedGraphToBigWig (v4) from kentUtils (https://github.com/ucscGenomeBrowser/kent). For the DSBCapture data, bedGraph files containing DSB locations were directly obtained from the Gene Expresion Omnibus (GSE78172) and converted to bigWig files using bedGraphToBigWig (v4). Three biological replicate samples (SRR7346494, SRR7346495, and SRR7346496) were summed for the sBLISS data, and two biological replicate samples (GSM2068755 and GSM2068756) were averaged for the DSBcapture data, in accordance with previously published methods [43]. DSB counts per 510 token window were then calculated by using bigWigAverageOverBed (v4) from kenUtils (https://github.com/ucscGenomeBrowser/kent) which calculates the average number of DSBCapture reads or sBLISS UMIs per base pair, therefore normalizing for the differences in window lengths. Genomic windows overlapping centromeric or telomeric regions (downloaded from the UCSC genome browser, https://www.genome.ucsc.edu/), or difficult to map regions (ENCODE blacklist version 1 [66]) were removed from the dataset using BEDtools intersect [67] (v2.26.0). Finally, genomic windows with DSB counts greater than the 99.99^th^ percentile were removed from the dataset to stabilize model training.

### GROVER fine-tuning

Genomic windows were split into train, validation and test sets using an 80:10:10 split for model training. All performance metrics shown were calculated using the unseen test set alone. DSB counts were scaled using StandardScaler from scikit-learn (v1.2.2) before training and testing. This scaling was later reversed for plotting. The pre-trained GROVER model was downloaded from HuggingFace (https://huggingface.co/PoetschLab/GROVER) using huggingface_hub (v0.20.3) and fine-tuned using Pytorch (v3.11.5) with an MSE loss function, using the AdamW optimizer with an epsilon of 10^-8^, a learning rate of 10^-6^, no warmup, a batch size of 32, and a dropout probability of 0.2. Fine-tuning hyperparameters were optimized using a randomized gird search on 1% of the dataset. The models were trained for 5 epochs but the performance on the validation set was assessed after each epoch and only the best performing model was kept.

### Density plots

Density plots were created using R (v4.4.0) with ggplot2 (v3.5.1). The densities were calculated using the stat_density_2d function of ggplot2, with n = 1000 and adjust = 1.5. A random sample of 10% of the dataset is shown in the density plots.

### Genome biological features

Chromatin mark and regulatory feature datasets from ENCODE for MCF7 and NHEK cells were obtained for the hg19 genome from UCSC genome browser in the form of bigWig files. RepliSeq data was downloaded from the UWRepliSeq dataset (http://hgdownload.cse.ucsc.edu/goldenpath/hg19/encodeDCC/wgEncodeUwRepliSeq) and DNAse-seq data was downloaded from the OpenChromDnase dataset (http://hgdownload.cse.ucsc.edu/goldenpath/hg19/encodeDCC/wgEncodeOpenChromDnas <u>e</u>). RNAseq data for MCF7 cells was downloaded from the CaltechRnaseq dataset (http://hgdownload.cse.ucsc.edu/goldenpath/hg19/encodeDCC/wgEncodeCaltechRnaSeq) and RNAseq data for NHEK cells was downloaded from the RegTxn dataset (http://hgdownload.cse.ucsc.edu/goldenpath/hg19/encodeDCC/wgEncodeRegTxn). The remaining datasets for NHEK were downloaded from the BroadHistone dataset (http://hgdownload.cse.ucsc.edu/goldenpath/hg19/encodeDCC/wgEncodeBroadHistone). H3K9me3, H3K27ac, H3K27me3, H3K36me3, and input data for MCF7 cells was downloaded from the SydhHistone dataset (http://hgdownload.cse.ucsc.edu/goldenpath/hg19/encodeDCC/wgEncodeSydhHistone). H3K4me3 data for MCF7 was downloaded from the UwHistone dataset (http://hgdownload.cse.ucsc.edu/goldenpath/hg19/encodeDCC/wgEncodeUwHistone). Pol2 and CTCF datasets for MC7 cells were downloaded from the OpenChromChip dataset (http://hgdownload.cse.ucsc.edu/goldenpath/hg19/encodeDCC/wgEncodeOpenChromChip). BigWig files were then averaged over the genomic windows using bigWigAverageOverBed (v4) from kenUtils (https://github.com/ucscGenomeBrowser/kent) to obtain a table of chromatin features per genomic window.

### Genome biological feature model and feature importances

Genomic windows were split into train, validation and test sets using an 80:10:10 split for model training. All performance metrics shown were calculated using the unseen test set alone. Random forest, XGBoost, LightGBM, MLP, and linear regression models were trained to predict DSB counts per window using the genome biological features for each window. DSB counts were scaled using StandardScaler from scikit-learn (v1.2.2) before training and testing. This scaling was later reversed for plotting. Genome biological features were also scaled in the same way for model training. The models were trained with scikit-learn (v1.2.2) using 500 estimators with no minimum samples per split or per leaf. For all models but linear regression (due to inferior performance), feature importances were calculated by training 10 models, measuring their performance using MSE, measuring the decrease in MSE when each feature is shufled across sequences, and then averaging the decreases in MSE across the 10 models. As an alternative approach, we also removed each feature one by one from the random forest model and measured the decrease in MSE of the model when each feature is removed. This process was also repeated 10 times and the average decrease in MSE was reported for each feature.

### Genome annotations

The gene and SINE annotation files for the hg19 genome were obtained from the UCSC genome browser (https://www.genome.ucsc.edu/). Promoter annotations were generated from the chromatin colors for NHEK cells (downloaded from the UCSC genome browser) and for MCF7 cells (downloaded from the Gene Expression Omnibus, GSE57498).

### Chromatin feature model with GROVER predictions

To incorporate GROVER predictions into the random forest model, we fine-tuned GROVER on two-thirds of the data in each fold and used it to generate predictions for the remaining one-third, following the standard three-fold cross-validation procedure. This process was repeated across all folds to produce a single prediction for each sequence in the entire dataset without redundancy. These predictions were then included within the table of chromatin features per window, and this table was then used to train the random forest models as before.

### Dimensionality reduction and GROVER embeddings

UMAP was performed using the uwot package (v0.2.2) in R (v4.4.0) with 15 neighbors, a minimum distance of 0.25 and the Euclidian distance metric. PCA was performed with centering and scaling using the stats package (v4.4.0) in R (v4.4.0). The data for the UMAP of pre-trained GROVER embeddings was obtained by using the 510-token genomic windows as input into the pre-trained GROVER model to obtain the 512 by 768-dimension matrices that make up the outputs of the final transformer layer of the model for each sequence. The embeddings from the first and last tokens, which are special tokens added to the sequence by the model, were removed from these matrices, and the embeddings were averaged across all tokens to obtain one 768-dimensional vector per genomic window which was then used for dimensionality reduction. To perform these analyses on the chromatin data, the tables of chromatin features for each genomic window that were used in the chromatin feature random forest models were also used for dimensionality reduction.

### GROVER model with integrated chromatin features

The fine-tuned GROVER model’s default classifier head was replaced with a neural network with one hidden layer containing 10000 nodes and a number of input nodes equal to 768 (length of the output of the pre-trained GROVER model) plus one extra input node per chromatin feature. These models were fine-tuned as before except this time with a learning rate of 10^-5^. As previously described, the hyperparameters were optimized using a randomized gird search on 1% of the dataset.

### Statistics

Correlation coefficients were calculated in R (v4.4.0) using the cor function of the stats package (v4.4.0). Wilcoxon rank-sum test was also calculated in R (v4.4.0) using the wilcox.test function of the stats package (v4.4.0).

## Declarations

### Ethics approval and consent to participate

Not applicable.

### Consent for publication

Not applicable.

### Availability of data and materials

All scripts, input and analysis files are available on Zenodo (https://doi.org/10.5281/zenodo.16375950).

### Author contributions

**ARP** conceptualized the study. **PMJ** and **MS** trained the models. **PMJ, MS,** and **ARP** analyzed the data. **PMJ** and **ARP** wrote the manuscript.

## Acknowledgements

This work was supported by the Center for Scalable data analytics and artificial intelligence (Scads.AI) Dresden-Leipzig. This work was partially funded by the Center for Advanced Systems Understanding (CASUS) which is financed by Germany’s Federal Ministry of Education and Research (BMBF) and by the Saxon Ministry for Science, Culture, and Tourism (SMWK) with tax funds on the basis of the budget approved by the Saxon State Parliament.

**ARP** was supported by the Mildred Scheel Early Career Center Dresden P2, funded by the German Cancer Aid. The authors gratefully acknowledge the computing time made available to them on the high-performance computer at the NHR Center of TU Dresden. This center is jointly supported by the Federal Ministry of Education and Research and the state governments participating in the NHR (www.nhr-verein.de/unsere-partner). We thank Jakub Zastapilo and Mario Aguilar-Herrador for their feedback on data visualization, and the Biomedical Genomics group at TU Dresden for helpful discussions throughout the research process.

## Additional files

### Additional file 1

**Fig. S1.** Validation of GROVER importance in combined double strand break prediction model. **Fig. S2.** Double strand break counts for promoters, stratified by GC content. **Fig. S3.** Uniform manifold approximation and projection (UMAP) analysis of chromatin features. **Fig. S4.** Performance and importances of models that predict double strand breaks (DSBs) within 50kbp genomic windows. **Fig. S5.** Performance and importances of models including ChIP-Seq input, trained to predict double strand breaks (DSBs). **Fig. S6.** Performance and importances of models using XGBoost. **Fig. S7.** Performance and importances of models using MLP. **Fig. S8.** Performance and importances of models using LightGBM. **Fig. S9.** Performance of models using Linear Regression. **Fig. S10.** Performance benchmarking.

**Fig. S1.**
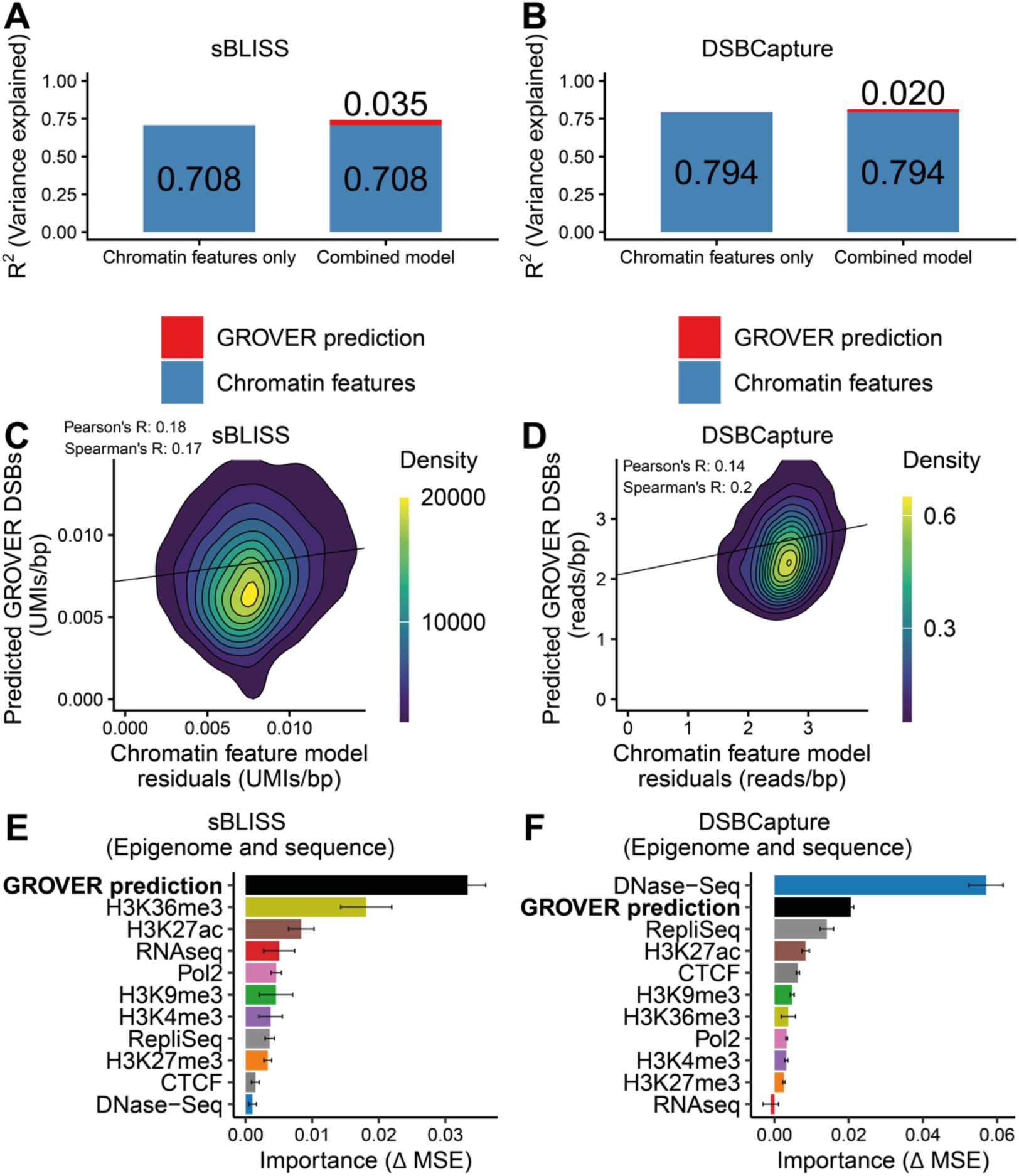
Validation of GROVER importance in combine double strand break prediction model. **A and B.** Variance explained by chromatin features and GROVER predictions in the combined double strand break prediction model for sBLISS data (**A**) and DSBCapture data (**B**). **C and D.** Correlation between chromatin feature model residuals and GROVER predictions for sBLISS data (**C**) and DSBCapture data (**D**). **E and F.** Feature ablation importances for combined double strand break prediction model for sBLISS data (**E**) and DSBCapture data (**F**). Importances are measured using the change in mean squared error (MSE) of the model when a feature is removed from the data and model. DSB data is shown as molecular identifier (UMI) counts (sBLISS) or read counts (DSBCapture) averaged per base pair over the length of each window.

**Fig. S2.**
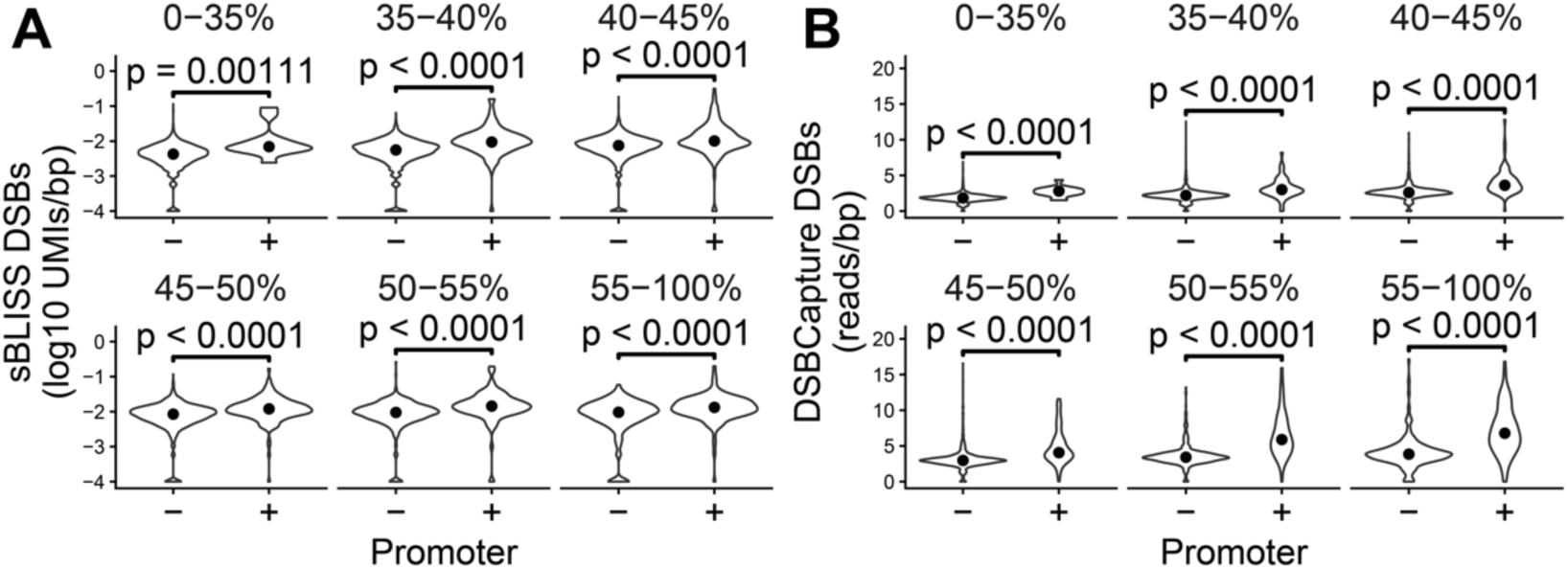
Double strand break counts for promoters, stratified by GC content. **A and B.** Double strand break counts for promoters according to sBLISS data (**A**) and DSBCapture data (**B**).

**Fig. S3.**
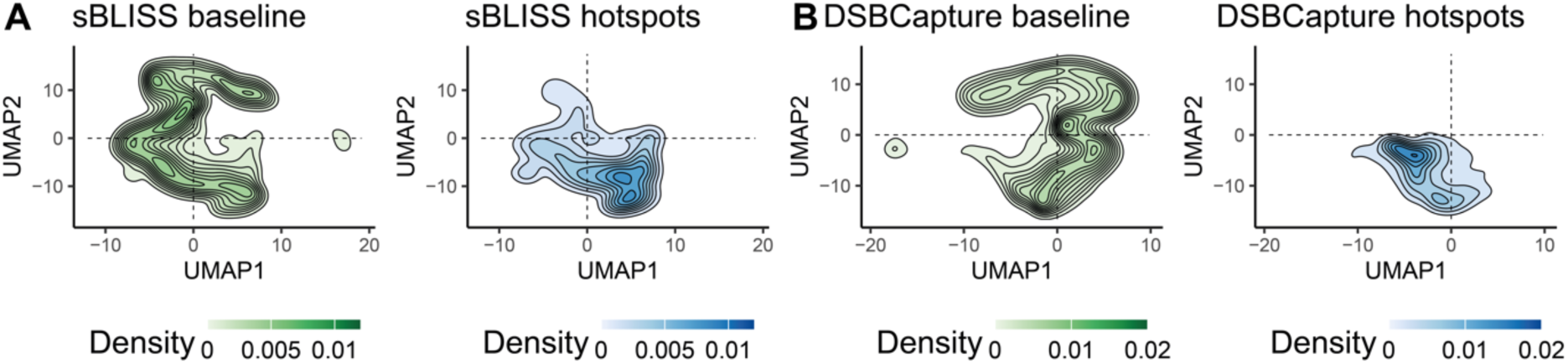
Uniform manifold approximation and projection (UMAP) analysis of chromatin features. **A and B.** UMAP of MCF7 (**A**) and NHEK (**B**) chromatin features. Sequences are separated based on whether they are in the top 10% (hotspots) or bottom 90% (baseline) of sequences.

**Fig. S4.**
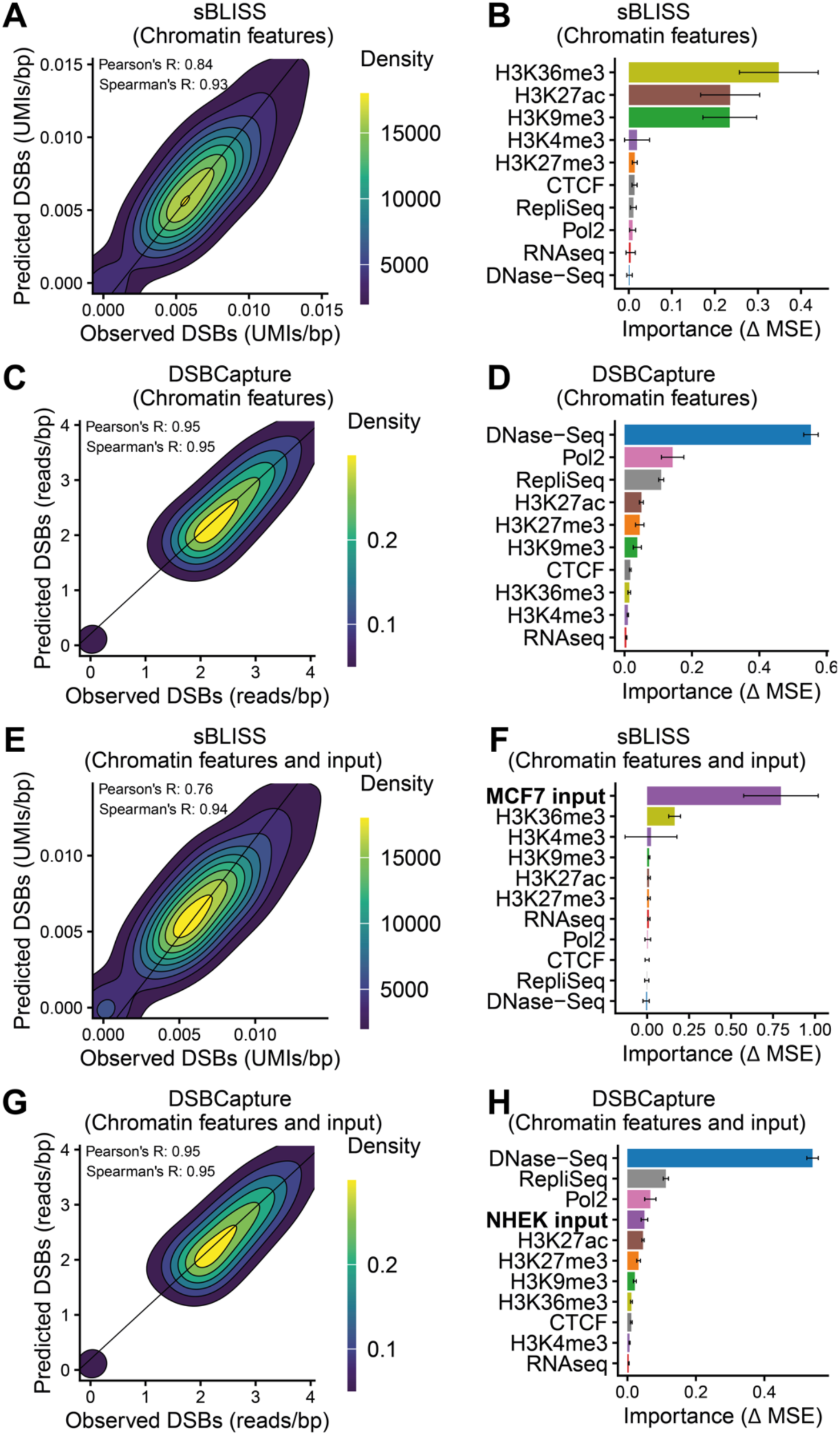
Performance and importances of models that predict double strand breaks (DSBs) within 50kbp genomic windows. **A and C.** Performance of random forest model trained on chromatin features to predict sBLISS data (**A**) and DSBCapture data (**C**). **B and D.** Importances of features used by the random forest model trained on chromatin features to predict sBLISS data (**B**) and DSBCapture data (**D**). **E and G.** Performance of random forest model trained on chromatin features and ChIP-Seq input data to predict sBLISS data (**E**) and DSBCapture data (**G**). **F and H.** Importances of features used by the random forest model trained on chromatin features and ChIP-Seq input data to predict sBLISS data (**F**) and DSBCapture data (**H**). DSB data is shown as molecular identifier (UMI) counts (sBLISS) or read counts (DSBCapture) averaged per base pair over the length of each window. Error bars represent standard deviation from the mean. Importances are measured using the change in mean squared error (MSE) of the model when a feature is shuffled in the test data.

**Fig. S5.**
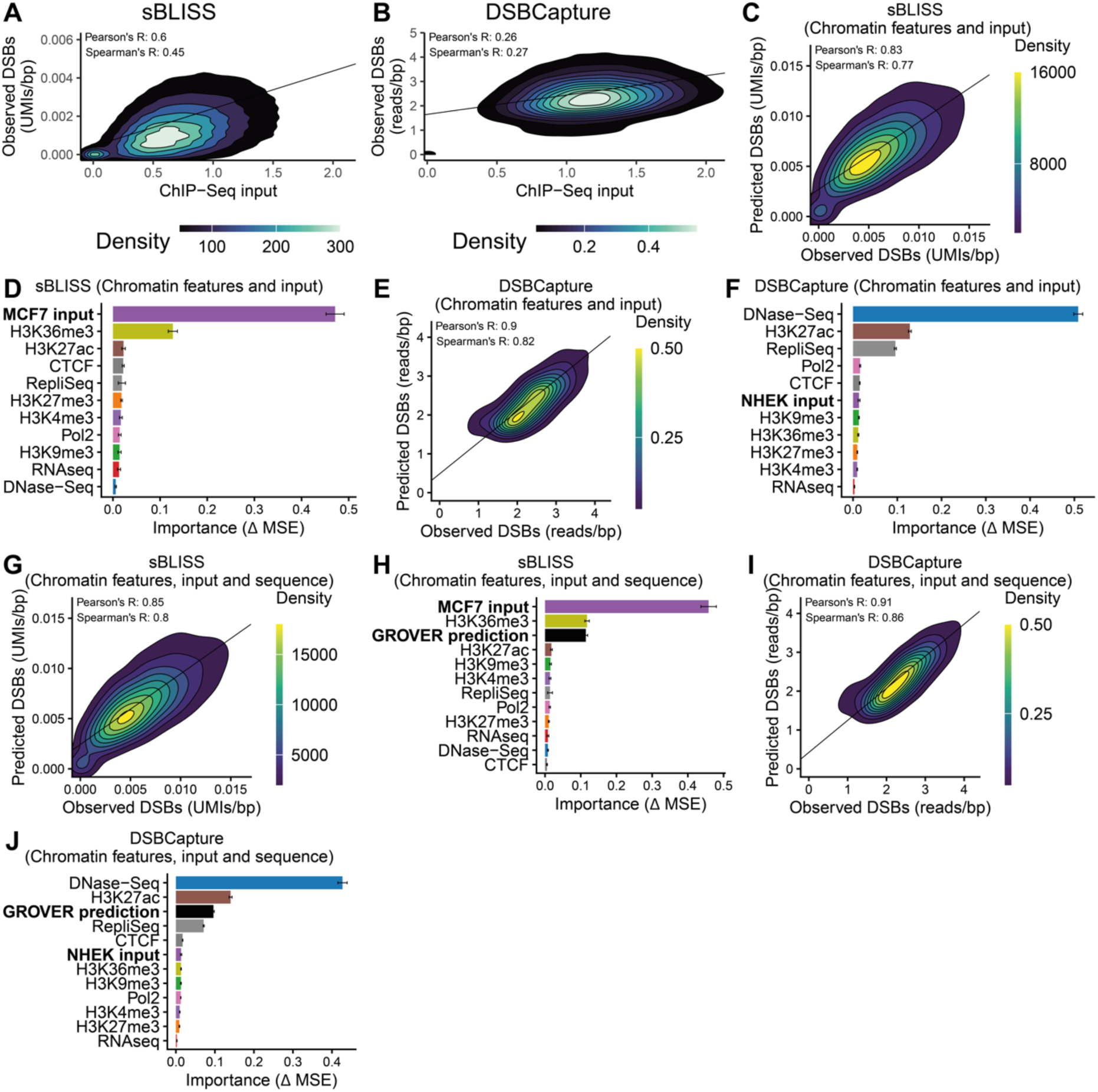
Performance and importances of models including ChIP-Seq input, trained to predict double strand breaks (DSBs). **A and B.** Correlation between ChIP-Seq input data and observed sBLISS data (**A**) or DSBCapture data (**B**). **C and E.** Performance of the random forest model trained on chromatin features and ChIP-Seq input data to predict sBLISS data (**C**) and DSBCapture data (**E**). **D and F.** Importances of features used by the random forest model trained on chromatin features and ChIP-Seq input data to predict sBLISS data (**D**) and DSBCapture data (**F**). **G and I.** Performance of the random forest model trained on chromatin features, ChIP-Seq input data and fine-tuned GROVER model predictions to predict sBLISS data (**G**) and DSBCapture data (**I**). **H and J.** Importances of features used by the random forest model trained on chromatin features, ChIP-Seq input data and fine-tuned GROVER model predictions to predict sBLISS data (**H**) and DSBCapture data (**J**). DSB data is shown as molecular identifier (UMI) counts (sBLISS) or read counts (DSBCapture) averaged per base pair over the length of each window. Error bars represent standard deviation from the mean. Importances are measured using the change in mean squared error (MSE) of the model when a feature is shuffled in the test data.

**Fig. S6.**
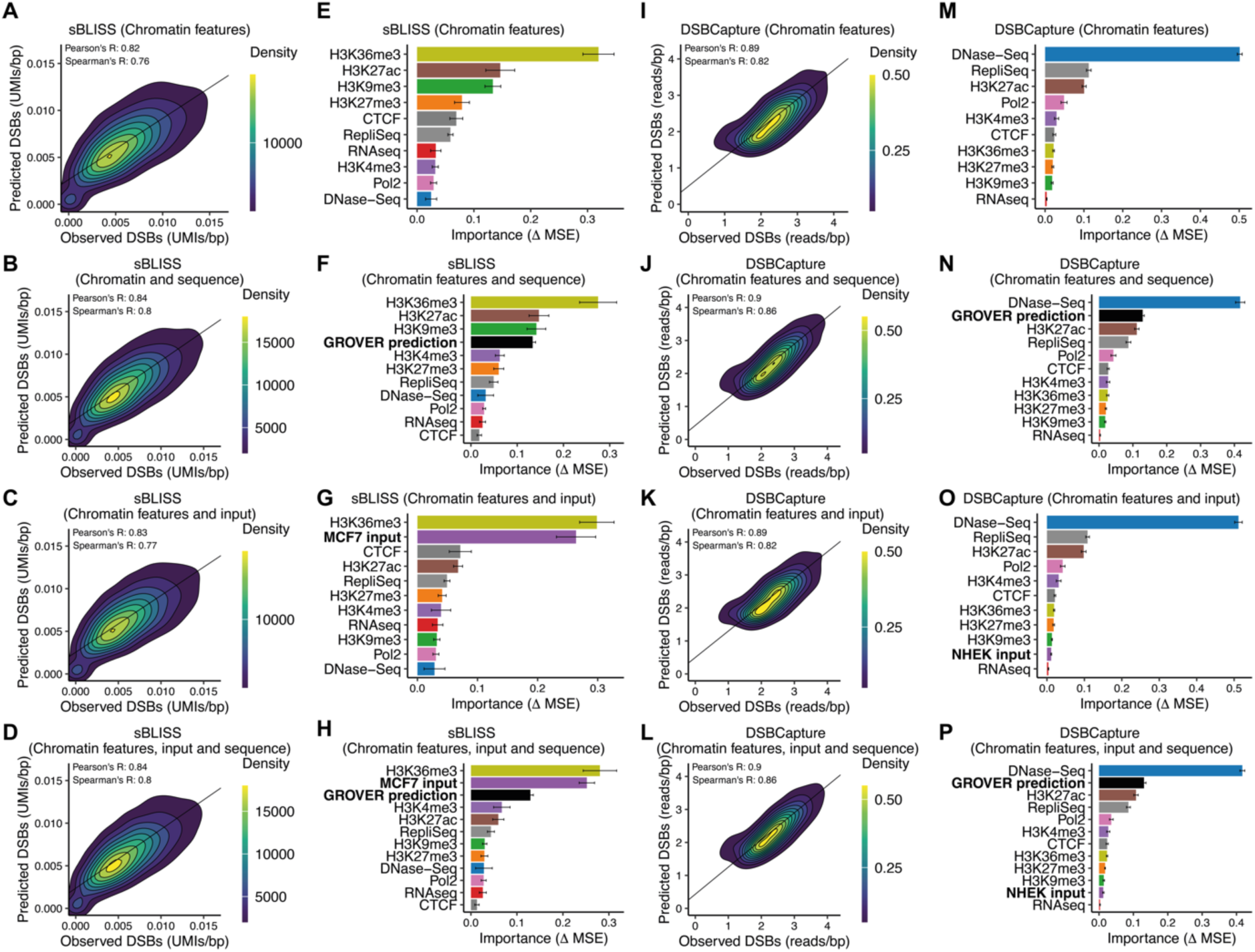
Performance and importances of models using eXtreme Gradient Boosting (XGBoost). **A.** Performance of XGBoost model to predict sBLISS data in MCF7 cells based on genome biological features, (**B)** genome biological features and GROVER, (**C)** genome biological features and ChIP-Seq input, and (**D)** genome biological features and ChIP-Seq input and GROVER. **E.** Feature contribution of XGBoost model to predict sBLISS data based on genome biological features, (**F**) genome biological features and GROVER, (**G**) genome biological features and ChIP-Seq input, and (**H**) genome biological features and ChIP-Seq input and GROVER. **I.** Performance of XGBoost model to predict DSBCapture data in NHEK cells based on genome biological features, **(J)** genome biological features and GROVER, **(K)** genome biological features and ChIP-Seq input, and **(L)** genome biological features and ChIP-Seq input and GROVER. **M.** Feature contribution of XGBoost model to predict sBLISS data based on genome biological features, **(N)** genome biological features and GROVER, **(O)** genome biological features and ChIP-Seq input, and **(P)** genome biological features and ChIP-Seq input and GROVER.

**Fig. S7.**
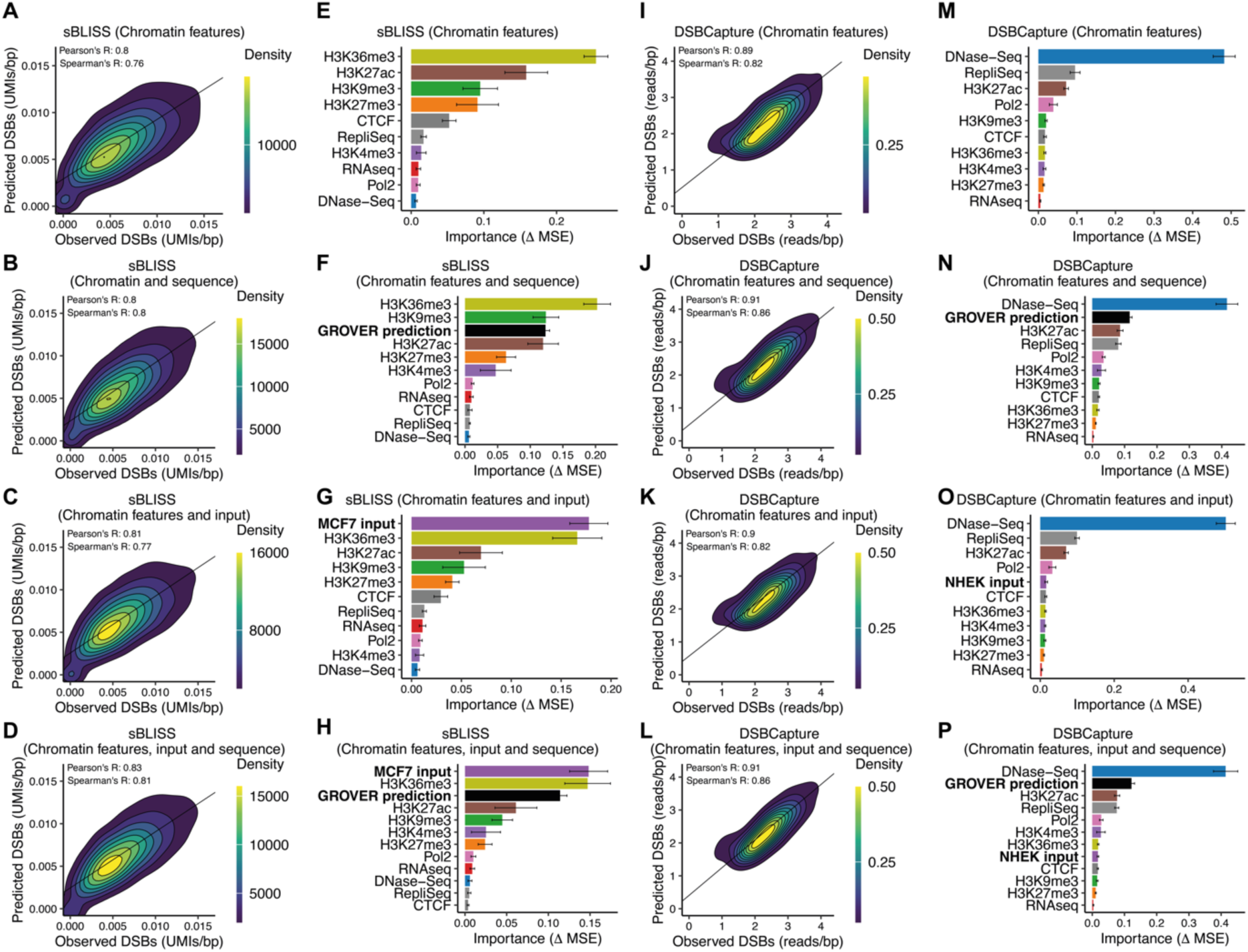
Performance and importances of models using Multilayer Perceptrons (MLP). **A.** Performance of MLP model to predict sBLISS data in MCF7 cells based on genome biological features, (**B)** genome biological features and GROVER, (**C)** genome biological features and ChIP-Seq input, and (**D)** genome biological features and ChIP- Seq input and GROVER. **E.** Feature contribution of XGBoost model to predict sBLISS data based on genome biological features, (**F**) genome biological features and GROVER, (**G**) genome biological features and ChIP-Seq input, and (**H**) genome biological features and ChIP-Seq input and GROVER. **I.** Performance of MLP model to predict DSBCapture data in NHEK cells based on genome biological features, **(J)** genome biological features and GROVER, **(K)** genome biological features and ChIP-Seq input, and **(L)** genome biological features and ChIP-Seq input and GROVER. **M.** Feature contribution of XGBoost model to predict sBLISS data based on genome biological features, **(N)** genome biological features and GROVER, **(O)** genome biological features and ChIP-Seq input, and **(P)** genome biological features and ChIP-Seq input and GROVER.

**Fig. S8.**
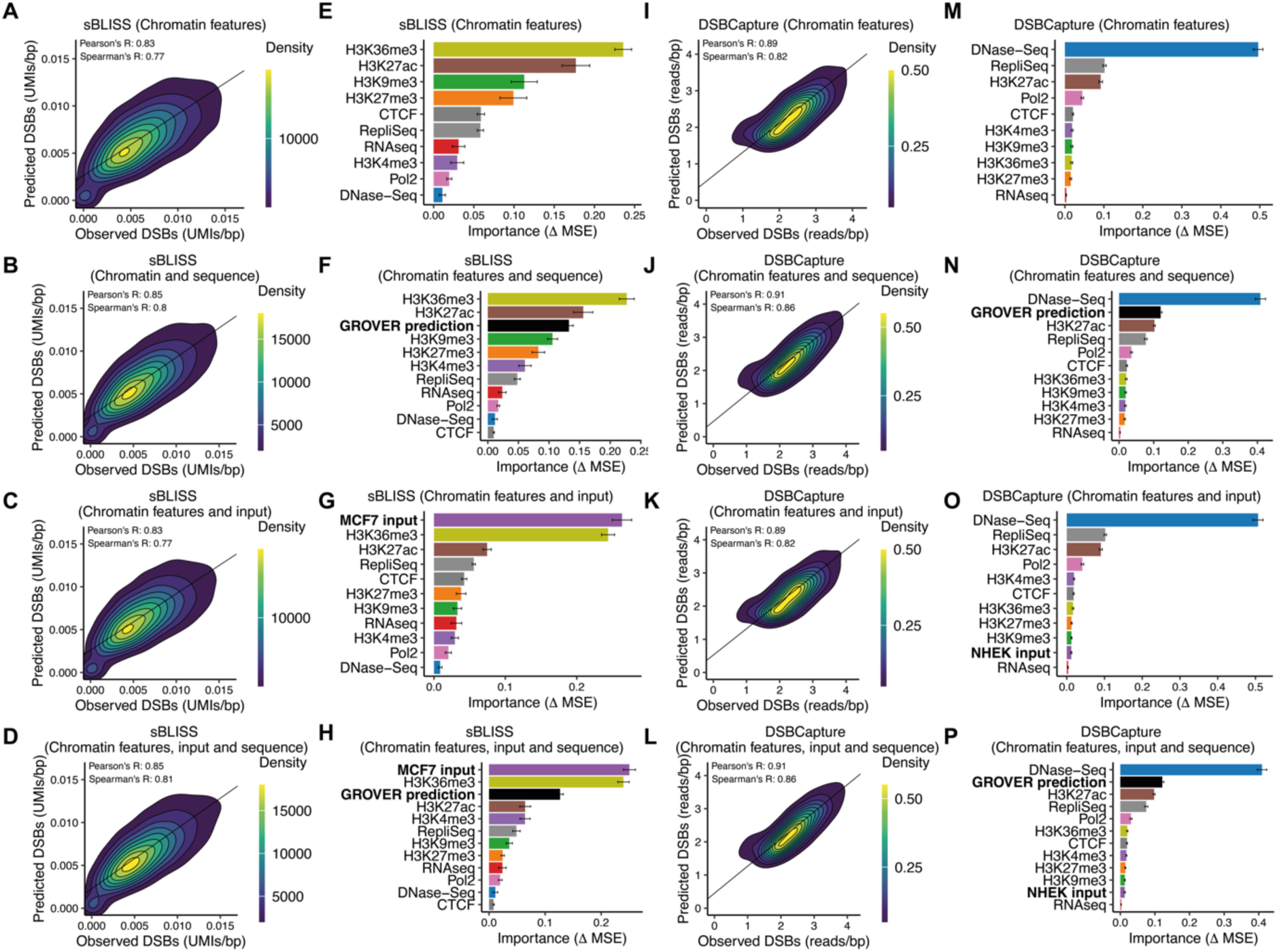
Performance and importances of models using Light Gradient Boosting Machine (LightGBM). **A.** Performance of LightGBM model to predict sBLISS data in MCF7 cells based on genome biological features, (**B)** genome biological features and GROVER, (**C)** genome biological features and ChIP-Seq input, and (**D)** genome biological features and ChIP-Seq input and GROVER. **E.** Feature contribution of XGBoost model to predict sBLISS data based on genome biological features, (**F**) genome biological features and GROVER, (**G**) genome biological features and ChIP-Seq input, and (**H**) genome biological features and ChIP-Seq input and GROVER. **I.** Performance of LightGBM model to predict DSBCapture data in NHEK cells based on genome biological features, **(J)** genome biological features and GROVER, **(K)** genome biological features and ChIP-Seq input, and **(L)** genome biological features and ChIP- Seq input and GROVER. **M.** Feature contribution of XGBoost model to predict sBLISS data based on genome biological features, **(N)** genome biological features and GROVER, **(O)** genome biological features and ChIP-Seq input, and **(P)** genome biological features and ChIP-Seq input and GROVER.

**Fig. S9.**
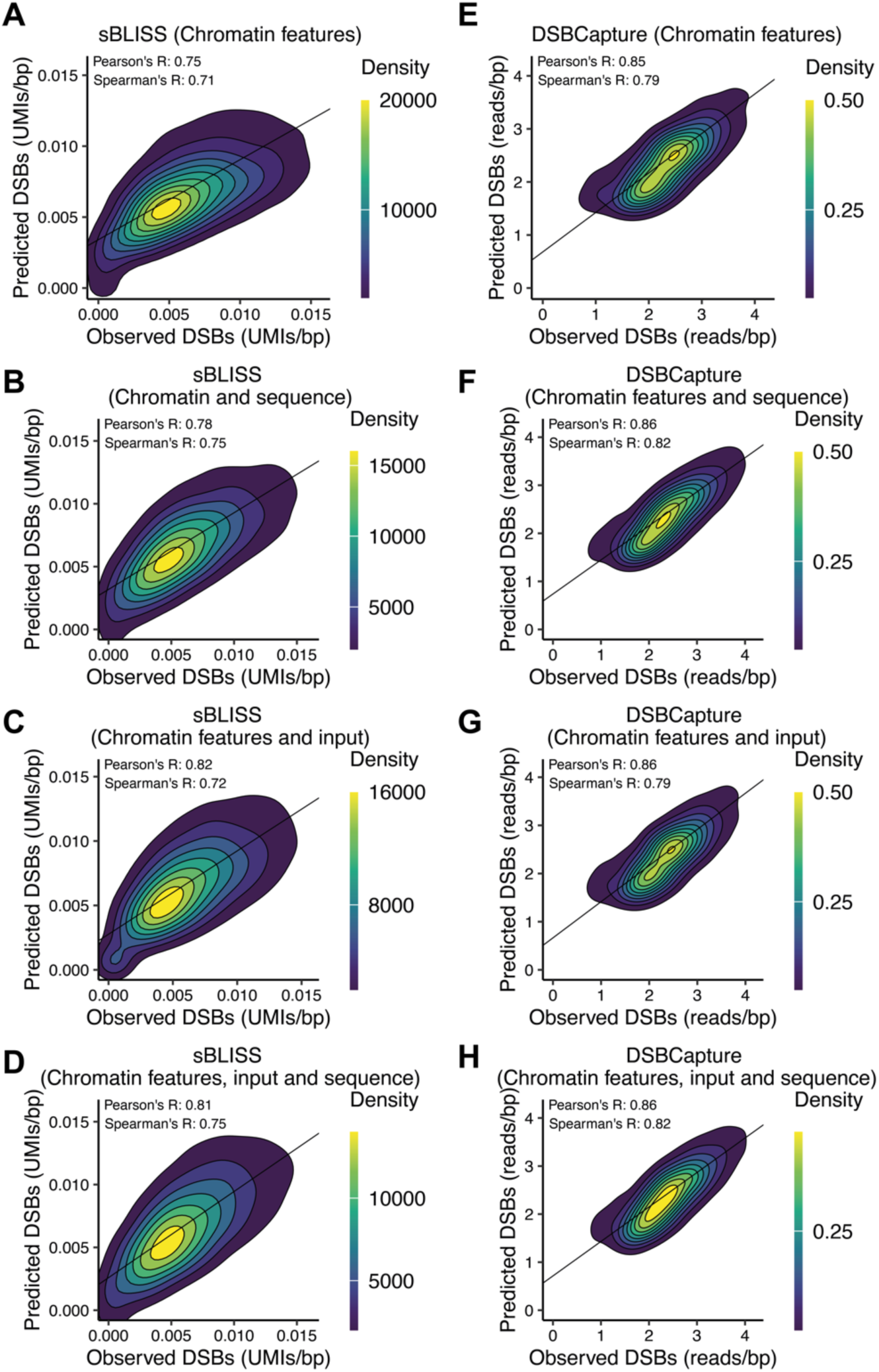
Performance of models using Linear Regression. **A.** Performance of a Linear Regression model to predict sBLISS data in MCF7 cells based on genome biological features, (**B)** genome biological features and GROVER, (**C)** genome biological features and ChIP-Seq input, and (**D)** genome biological features and ChIP-Seq input and GROVER.

**Fig. S10.**
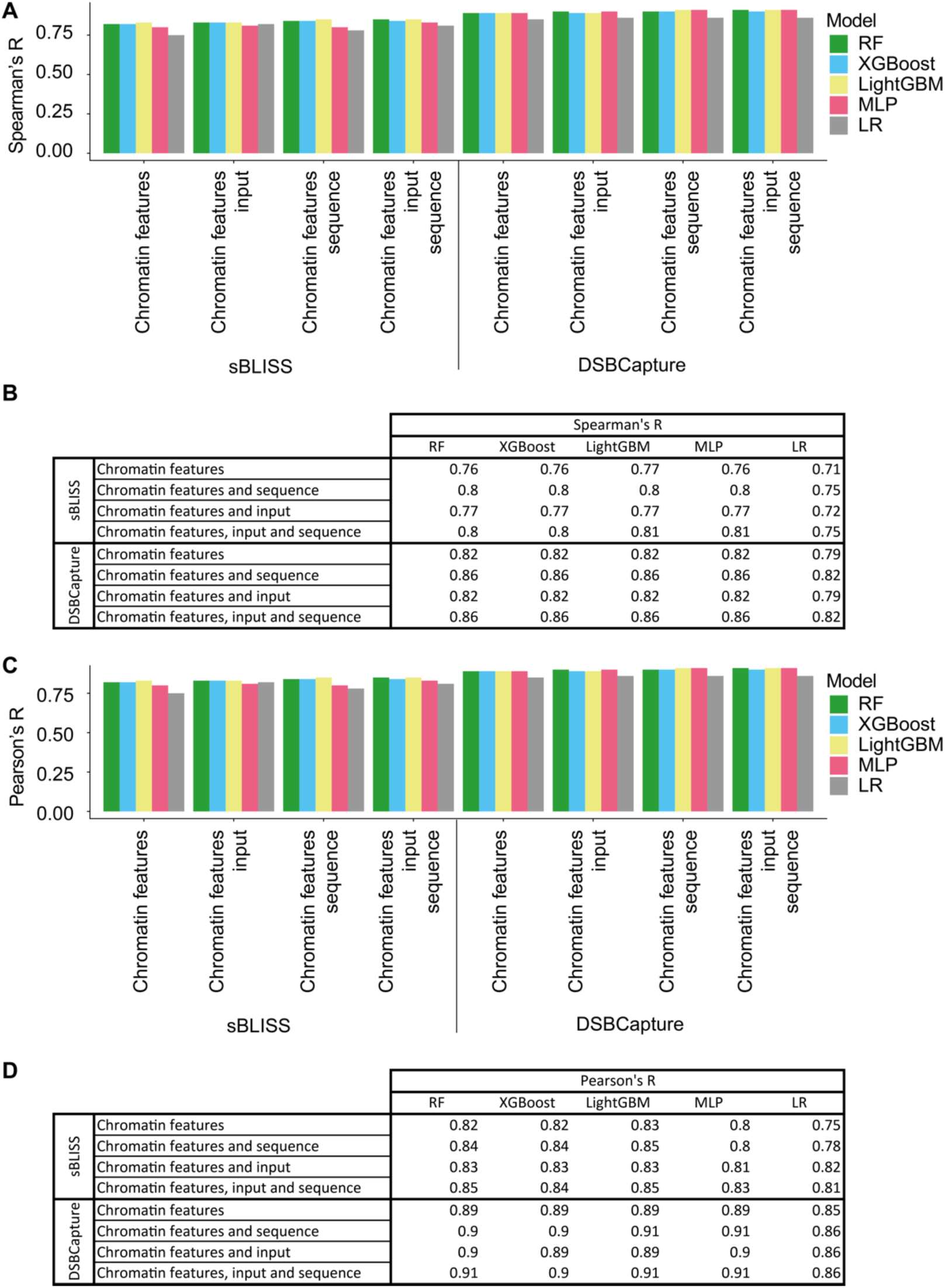
Performance benchmarking. **A & B.** Spearman’s R for Random Forest (RF), eXtreme Gradient Boosting (XGBoost), Light Gradient Boosting Machine (LightGBM), Multy Layer Perceptron (MLP), and Linear Regression (LR) for different sets of input data to predict sBLISS in MCF7 cells and DSBCapture in NHEK cells. **C & D.** Pearson’s R for Random Forest (RF), eXtreme Gradient Boosting (XGBoost), Light Gradient Boosting Machine (LightGBM), Multy Layer Perceptron (MLP), and Linear Regression (LR) for different sets of input data to predict sBLISS in MCF7 cells and DSBCapture in NHEK cells.

## References

1. Pommier Y, Nussenzweig A, Takeda S, Austin C. Human topoisomerases and their roles in genome stability and organization. Nature Reviews Molecular Cell Biology. 2022 June 1;23(6):407–27.

2. Castillo-Guzman D, Chédin F. Defining R-loop classes and their contributions to genome instability. DNA Repair. 2021 Oct 1;106:103182.

3. Crossley MP, Bocek M, Cimprich KA. R-Loops as Cellular Regulators and Genomic Threats. Molecular Cell. 2019 Feb 7;73(3):398–411.

4. Puget N, Miller KM, Legube G. Non-canonical DNA/RNA structures during Transcription-Coupled Double-Strand Break Repair: Roadblocks or Bona fide repair intermediates? DNA Repair. 2019 Sept 1;81:102661.

5. Técher H, Koundrioukoff S, Nicolas A, Debatisse M. The impact of replication stress on replication dynamics and DNA damage in vertebrate cells. Nature Reviews Genetics. 2017 Sept 1;18(9):535–50.

6. Kotsantis P, Segura-Bayona S, Margalef P, Marzec P, Ruis P, Hewitt G, et al. RTEL1 Regulates G4/R-Loops to Avert Replication-Transcription Collisions. Cell Rep. 2020 Dec 22;33(12):108546.

7. Larsen F, Gundersen G FAU - Lopez R, Lopez R FAU - Prydz H, Prydz H, Kruisselbrink E, Guryev V FAU - Brouwer K, et al. CpG islands as gene markers in the human genome. (0888-7543 (Print)).

8. Lander ES, Linton LM, Birren B, Nusbaum C, Zody MC, Baldwin J, et al. Initial sequencing and analysis of the human genome. Nature. 2001 Feb 1;409(6822):860–921.

9. Crosetto N, Mitra A, Silva MJ, Bienko M, Dojer N, Wang Q, et al. Nucleotide-resolution DNA double-strand break mapping by next-generation sequencing. Nature Methods. 2013 Apr 1;10(4):361–5.

10. Morales ME, White TB, Streva VA, DeFreece CB, Hedges DJ, Deininger PL. The Contribution of Alu Elements to Mutagenic DNA Double-Strand Break Repair. PLOS Genetics. 2015 Mar 11;11(3):e1005016.

11. Lehrman MA, Schneider WJ, Südhof TC, Brown MS, Goldstein JL, Russell DW. Mutation in LDL Receptor: Alu-Alu Recombination Deletes Exons Encoding Transmembrane and Cytoplasmic Domains. Science. 1985 Jan 11;227(4683):140–6.

12. Gasior SL, Wakeman TP, Xu B, Deininger PL. The Human LINE-1 Retrotransposon Creates DNA Double-strand Breaks. Journal of Molecular Biology. 2006 Apr 14;357(5):1383–93.

13. Gadgil RY, Romer EJ, Goodman CC, Rider SD, Damewood FJ, Barthelemy JR, et al. Replication stress at microsatellites causes DNA double-strand breaks and break-induced replication. Journal of Biological Chemistry. 2020 Nov 6;295(45):15378–97.

14. Neil AJ, Kim JC, Mirkin SM. Precarious maintenance of simple DNA repeats in eukaryotes. Bioessays. 2017 Sept;39(9).

15. Correll-Tash S, Lilley B, Salmons IV H, Mlynarski E, Franconi CP, McNamara M, et al. Double strand breaks (DSBs) as indicators of genomic instability in PATRR-mediated translocations. Human Molecular Genetics. 2020 Dec 15;29(24):3872–81.

16. Glover TW, Wilson TE, Arlt MF. Fragile sites in cancer: more than meets the eye. Nature Reviews Cancer. 2017 Aug 1;17(8):489–501.

17. Chakrabarti AM, Henser-Brownhill T, Monserrat J, Poetsch AR, Luscombe NM, Scaffidi P. Target-Specific Precision of CRISPR-Mediated Genome Editing. Molecular Cell. 2019 Feb 21;73(4):699–713.e6.

18. Shou J, Li J, Liu Y, Wu Q. Precise and Predictable CRISPR Chromosomal Rearrangements Reveal Principles of Cas9-Mediated Nucleotide Insertion. Molecular Cell. 2018 Aug 16;71(4):498–509.e4.

19. Shen MW, Arbab M, Hsu JY, Worstell D, Culbertson SJ, Krabbe O, et al. Predictable and precise template-free CRISPR editing of pathogenic variants. Nature. 2018 Nov;563(7733):646–51.

20. Milano L, Gautam A, Caldecott KW. DNA damage and transcription stress. Molecular Cell. 2024 Jan 4;84(1):70–9.

21. Gothe HJ, Bouwman BAM, Gusmao EG, Piccinno R, Petrosino G, Sayols S, et al. Spatial Chromosome Folding and Active Transcription Drive DNA Fragility and Formation of Oncogenic MLL Translocations. Molecular Cell. 2019 July 25;75(2):267–283.e12.

22. Creyghton MP, Cheng AW, Welstead GG, Kooistra T, Carey BW, Steine EJ, et al. Histone H3K27ac separates active from poised enhancers and predicts developmental state. Proceedings of the National Academy of Sciences. 2010 Dec 14;107(50):21931–6.

23. Heintzman ND, Hon GC, Hawkins RD, Kheradpour P, Stark A, Harp LF, et al. Histone modifications at human enhancers reflect global cell-type-specific gene expression. Nature. 2009 May 1;459(7243):108–12.

24. Duttke SHC, Lacadie SA, Ibrahim MM, Glass CK, Corcoran DL, Benner C, et al. Human Promoters Are Intrinsically Directional. Molecular Cell. 2015 Feb 19;57(4):674–84.

25. Wang H, Helin K. Roles of H3K4 methylation in biology and disease. Trends in Cell Biology. 2025 Feb 1;35(2):115–28.

26. Sharda A, Humphrey TC. The role of histone H3K36me3 writers, readers and erasers in maintaining genome stability. DNA Repair. 2022 Nov 1;119:103407.

27. Li Z, Zhang Z. The silent guardian: unraveling the roles of H3K9me3 in genome maintenance. Genome Instability & Disease. 2024 Aug 1;5(4):133–53.

28. Bernstein BE, Mikkelsen TS, Xie X, Kamal M, Huebert DJ, Cuff J, et al. A Bivalent Chromatin Structure Marks Key Developmental Genes in Embryonic Stem Cells. Cell. 2006 Apr 21;125(2):315–26.

29. Kundu S, Ji F, Sunwoo H, Jain G, Lee JT, Sadreyev RI, et al. Polycomb Repressive Complex 1 Generates Discrete Compacted Domains that Change during Differentiation. Molecular Cell. 2017 Feb 2;65(3):432–446.e5.

30. Hosogane M, Funayama R, Shirota M, Nakayama K. Lack of Transcription Triggers H3K27me3 Accumulation in the Gene Body. Cell Reports. 2016 July 19;16(3):696–706.

31. Ayrapetov MK, Gursoy-Yuzugullu O, Xu C, Xu Y, Price BD. DNA double-strand breaks promote methylation of histone H3 on lysine 9 and transient formation of repressive chromatin. Proceedings of the National Academy of Sciences. 2014 June 24;111(25):9169–74.

32. Zhang Y, Chang JF, Sun J, Chen L, Yang XM, Tang HY, et al. Histone H3K27 methylation modulates the dynamics of FANCD2 on chromatin to facilitate NHEJ and genome stability. Journal of Cell Science. 2018 June 21;131(12):jcs215525.

33. Hilmi K, Jangal M, Marques M, Zhao T, Saad A, Zhang C, et al. CTCF facilitates DNA double-strand break repair by enhancing homologous recombination repair. Science Advances. 3(5):e1601898.

34. Lang F, Li X, Zheng W, Li Z, Lu D, Chen G, et al. CTCF prevents genomic instability by promoting homologous recombination-directed DNA double-strand break repair. Proceedings of the National Academy of Sciences. 2017 Oct 10;114(41):10912–7.

35. Canela A, Maman Y, Jung S, Wong N, Callen E, Day A, et al. Genome Organization Drives Chromosome Fragility. Cell. 2017 July 27;170(3):507–521.e18.

36. Helmrich A, Ballarino M, Tora L. Collisions between Replication and Transcription Complexes Cause Common Fragile Site Instability at the Longest Human Genes. Molecular Cell. 2011 Dec 23;44(6):966–77.

37. Wilson TE, Arlt MF, Park SH, Rajendran S, Paulsen M, Ljungman M, et al. Large transcription units unify copy number variants and common fragile sites arising under replication stress. Genome Res. 2015 Feb;25(2):189–200.

38. Barlow JH, Faryabi RB, Callén E, Wong N, Malhowski A, Chen HT, et al. Identification of early replicating fragile sites that contribute to genome instability. Cell. 2013 Jan 31;152(3):620–32.

39. Tubbs A, Sridharan S, van Wietmarschen N, Maman Y, Callen E, Stanlie A, et al. Dual Roles of Poly(dA:dT) Tracts in Replication Initiation and Fork Collapse. Cell. 2018 Aug 23;174(5):1127–1142.e19.

40. Lensing SV, Marsico G, Hänsel-Hertsch R, Lam EY, Tannahill D, Balasubramanian S. DSBCapture: in situ capture and sequencing of DNA breaks. Nat Methods. 2016 Oct;13(10):855–7.

41. Yan WX, Mirzazadeh R, Garnerone S, Scott D, Schneider MW, Kallas T, et al. BLISS is a versatile and quantitative method for genome-wide profiling of DNA double-strand breaks. Nat Commun. 2017 May 12;8(1):15058.

42. Bouwman BAM, Agostini F, Garnerone S, Petrosino G, Gothe HJ, Sayols S, et al. Genome-wide detection of DNA double-strand breaks by in-suspension BLISS. Nature Protocols. 2020 Dec 1;15(12):3894–941.

43. Ballinger TJ, Bouwman BAM, Mirzazadeh R, Garnerone S, Crosetto N, Semple CA. Modeling double strand break susceptibility to interrogate structural variation in cancer. Genome Biology. 2019 Feb 8;20(1):28.

44. Mourad R, Ginalski K, Legube G, Cuvier O. Predicting double-strand DNA breaks using epigenome marks or DNA at kilobase resolution. Genome Biology. 2018 Mar 15;19(1):34.

45. Schep R, Brinkman EK, Leemans C, Vergara X, van der Weide RH, Morris B, et al. Impact of chromatin context on Cas9-induced DNA double-strand break repair pathway balance. Molecular Cell. 2021 May 20;81(10):2216–2230.e10.

46. Vergara X, Manjón AG, de Haas M, Morris B, Schep R, Leemans C, et al. Widespread chromatin context-dependencies of DNA double-strand break repair proteins. Nature Communications. 2024 June 22;15(1):5334.

47. Schep R, Leemans C, Brinkman EK, van Schaik T, van Steensel B. Protocol: A Multiplexed Reporter Assay to Study Effects of Chromatin Context on DNA Double-Strand Break Repair. Frontiers in Genetics [Internet]. 2022;Volume 12-2021. Available from: https://www.frontiersin.org/journals/genetics/articles/10.3389/fgene.2021.785947

48. Vaswani A, Shazeer N, Parmar N, Uszkoreit J, Jones L, Gomez AN, et al. Attention Is All You Need. CoRR [Internet]. 2017;abs/1706.03762. Available from: http://arxiv.org/abs/1706.03762

49. Avsec Ž, Agarwal V, Visentin D, Ledsam JR, Grabska-Barwinska A, Taylor KR, et al. Effective gene expression prediction from sequence by integrating long-range interactions. Nature Methods. 2021 Oct 1;18(10):1196–203.

50. Consens ME, Li B, Poetsch AR, Gilbert S. Genomic language models could transform medicine but not yet. npj Digital Medicine. 2025 Apr 18;8(1):212.

51. Benegas G, Ye C, Albors C, Li JC, Song YS. Genomic language models: opportunities and challenges. Trends in Genetics. 2025 Apr 1;41(4):286–302.

52. Sanabria M, Hirsch J, Poetsch AR. Distinguishing word identity and sequence context in DNA language models. BMC Bioinformatics. 2024 Sept 13;25(1):301.

53. Sanabria M, Hirsch J, Joubert PM, Poetsch AR. DNA language model GROVER learns sequence context in the human genome. Nat Mach Intell. 2024 Aug;6(8):911–23.

54. Hashimshony T, Wagner F, Sher N, Yanai I. CEL-Seq: Single-Cell RNA-Seq by Multiplexed Linear Amplification. Cell Reports. 2012 Sept 27;2(3):666–73.

55. Van Gelder RN, von Zastrow ME, Yool A, Dement WC, Barchas JD, Eberwine JH. Amplified RNA synthesized from limited quantities of heterogeneous cDNA. Proceedings of the National Academy of Sciences. 1990 Mar 1;87(5):1663–7.

56. van Bueren MAE, Janssen A. The impact of chromatin on double-strand break repair: Imaging tools and discoveries. DNA Repair. 2024 Jan 1;133:103592.

57. Dunham I, Kundaje A, Aldred SF, Collins PJ, Davis CA, Doyle F, et al. An integrated encyclopedia of DNA elements in the human genome. Nature. 2012 Sept 1;489(7414):57– 74.

58. Beware Default Random Forest Importances [Internet]. [cited 2025 Jan 27]. Available from: http://explained.ai/decision-tree-viz/index.html

59. Chen B, Ren C, Ouyang Z, Xu J, Xu K, Li Y, et al. Stratifying TAD boundaries pinpoints focal genomic regions of regulation, damage, and repair. Briefings in Bioinformatics. 2024 July 1;25(4):bbae306.

60. Mao Z, Jiang Y, Liu X, Seluanov A, Gorbunova V. DNA repair by homologous recombination, but not by nonhomologous end joining, is elevated in breast cancer cells. Neoplasia. 2009 July;11(7):683–91.

61. UW Repli-seq Track Settings [Internet]. [cited 2025 Jan 27]. Available from: https://genome.ucsc.edu/cgi-bin/hgTrackUi?db=hg19&g=wgEncodeUwRepliSeq

62. Hansen RS, Thomas S, Sandstrom R, Canfield TK, Thurman RE, Weaver M, et al. Sequencing newly replicated DNA reveals widespread plasticity in human replication timing. Proceedings of the National Academy of Sciences. 2010 Jan 5;107(1):139–44.

63. Kasai H, Nishimura S. Hydroxylation of deoxyguanosine at the C-8 position by ascorbic acid and other reducing agents. Nucleic Acids Research. 1984 Feb 24;12(4):2137–45.

64. Li H. Aligning sequence reads, clone sequences and assembly contigs with BWA-MEM [Internet]. 2013. Available from: https://arxiv.org/abs/1303.3997

65. Li H, Handsaker B, Wysoker A, Fennell T, Ruan J, Homer N, et al. The Sequence Alignment/Map format and SAMtools. Bioinformatics. 2009 June;25(16):2078–9.

66. Amemiya HM, Kundaje A, Boyle AP. The ENCODE Blacklist: Identification of Problematic Regions of the Genome. Scientific Reports. 2019 June 27;9(1):9354.

67. Quinlan AR, Hall IM. BEDTools: a flexible suite of utilities for comparing genomic features. Bioinformatics. 2010 Jan;26(6):841–2.

68. Poetsch, A. R., Boulton, S. J. & Luscombe, N. M. Genomic landscape of oxidative DNA damage and repair reveals regioselective protection from mutagenesis. Genome Biol 19, 215 (2018).

69. Poetsch, A. R. The genomics of oxidative DNA damage, repair, and resulting mutagenesis. Comput Struct Biotechnol J 18, 207–219 (2020).

70. Takhaveev, V. et al. Click-code-seq reveals strand biases of DNA oxidation and depurination in human genome. Nature Chemical Biology 1–12 (2025).

71. Avsec, Ž., Latysheva, N., Cheng, J., et al. Advancing regulatory variant effect prediction with AlphaGenome. Nature 649, 1206–1218 (2026).

72. Ju, B-G. et al. A topoisomerase IIβ-mediated dsDNA break required for regulated transcription. Science 312, 1798–1802 (2006).

73. Verarga, X. et al. Widespread chromatin context-dependencies of DNA double-strand break repair proteins. Nature Communications 15 (2024).

